# Task-aligned foundation model representations enable single-cell cytokine response inference

**DOI:** 10.64898/2026.11.26.769597

**Authors:** Yujin L. Jeong, Jong Kyoung Kim

## Abstract

Inferring cellular responses from single-cell transcriptomes requires interpreting changes relative to an appropriate baseline while retaining cell-type context. Here we present CYTAP, which adapts frozen single-cell foundation model embeddings for cytokine-response inference by combining donor-adjusted receiver states with deviations from matched control centroids. CYTAP uses a shared classifier to assign relative single-cell response scores across 90 cytokines and supports unpaired analyses using counterfactual control references. Baseline-relative representations improved held-out-donor prediction with both State SE and scFoundation. Across independent perturbation datasets, CYTAP prioritized applied cytokines, including when receptor transcripts were sparsely detected, and ranked administered cytokines among the top ten in both in vivo datasets. In an unpaired COVID-19 cohort, inferred responses were associated with severity, and IL-6 response scores correlated with plasma IL-6 concentrations. These findings support the use of biologically defined references to adapt pretrained representations for functional response inference.

## Introduction

Inferring cellular responses from single-cell transcriptomes requires interpreting changes relative to an appropriate baseline while retaining the context of the responding cell. Single-cell foundation models provide reusable representations of transcriptomes, offering a common feature space for downstream analysis ^1–5^. For response inference, cell-type identity can provide essential context, whereas donor-associated baseline differences may complicate comparisons across individuals. Existing perturbation models and transfer-learning frameworks address related challenges involving treatment effects, covariates and reference mapping ^6–9^. A complementary strategy is to use biologically defined control references to construct task-aligned inputs from frozen embeddings, making baseline-relative changes explicit while preserving relevant cell-state context.

Cytokine-response inference provides a demanding application of this strategy. Cytokines are major mediators of intercellular communication in the immune system, and the same cytokine can induce distinct transcriptional programs in different receiver cell types^10,11^. Many methods for analyzing single-cell RNA sequencing (scRNA-seq) data infer such communication by linking ligand-expressing sender cells to cells expressing the corresponding receptor^12–15^. However, their predictions vary substantially with the interaction database and scoring algorithm^16,17^. Expression-based inference also faces two fundamental limitations. First, low detection rates of ligand and receptor transcripts can obscure candidate interactions^18–20^, even when the corresponding proteins are present^21^. Aggregating expression within predefined cell-type groups can mitigate sparsity, but constrains inference to the resolution of those groups. Second, transcript abundance alone does not establish whether signaling has occurred. Cytokine activity can depend on release or post-translational activation, as illustrated by the release of IL-1α and IL-33 following cell damage^22^, caspase-1-mediated processing of IL-1β and IL-18^23^, and integrin-mediated activation of latent TGF-β^24,25^. Moreover, receiver cells need not express every ligand-binding receptor subunit: IL-15 can be trans-presented by IL-15Rα to neighboring cells lacking IL-15Rα^26^, and IL-6 can signal through soluble IL-6R in cells lacking the membrane-bound IL-6R^27,28^.

Response-based methods address these limitations by relating candidate signals to downstream transcriptional changes in receiver cells. NicheNet^29^ links ligands to target genes, whereas CytoSig^30^ infers cytokine activity using response signatures derived from bulk transcriptomic profiles. IREA^10^ catalogues mouse immune responses to cytokines, SecAct^31^ infers secreted protein activity from spatial transcriptomes, and huCIRA^32^ draws on a human cytokine-response dictionary. ContactTracing^33^ evaluates receptor-associated responses by comparing receptor-expressing and receptor-null cells. These approaches differ in their use of cell-type-specific references, dependence on receptor detection and resolution of inference. Scupa applies a single-cell foundation model to quantify predefined polarization states rather than individual cytokine responses^34^. Together, these advances motivate a framework that assigns cytokine-response scores to individual cells while accounting for both the receiver-cell context and its deviation from an appropriate baseline.

Two recent developments provide an opportunity to address this challenge. First, the Parse Biosciences human peripheral blood mononuclear cell (PBMC) cytokine-stimulation atlas^35^ (Parse-PBMC) profiles approximately 9.7 million cells from 12 donors exposed to 90 cytokines, providing a large-scale reference of cell-type-dependent responses. Second, single-cell foundation models pretrained on large atlases provide cell-state representations that integrate transcriptome-wide information^1–5^. These embeddings offer a common feature space for comparing receiver states, but also contain baseline variation associated with cell identity and donor. Task-aligned adaptation of these representations to distinguish response-associated changes from baseline variation may therefore improve cytokine-response inference.

Here we present CYTAP (CYTokine response prediction by Adapting Pretrained embeddings), a framework for inferring cytokine responses in individual receiver cells from their transcriptional states. CYTAP uses embeddings from the pretrained State embedding (SE) model^1^ and combines a donor-adjusted receiver state with its deviation from the donor- and cell-type-matched control centroid. This composite representation retains receiver-cell context while explicitly representing baseline-relative changes. A shared classifier trained on Parse-PBMC assigns each cell relative response scores across the 90 cytokines. The representation builds on approaches that separate perturbation effects from covariates or align cell states in latent space^7–9^. When donor-matched controls are unavailable, pooled or CellOT-derived counterfactual control references enable application to unpaired datasets^6^.

Across independent in vitro and in vivo stimulation datasets, CYTAP prioritized the applied cytokines, including under conditions of sparse receptor transcript detection. CYTAP was the only evaluated method that placed the administered cytokine within the top ten candidates across both in vivo datasets. In an unpaired COVID-19 cohort, CYTAP identified severity-associated cytokine responses, including an IL-6 response score that correlated with independently measured plasma IL-6 concentrations. Collectively, these results support the use of baseline-relative receiver-cell states to infer cytokine-response programs without requiring detected ligand-receptor expression pairs, providing complementary evidence for cytokine-mediated communication analysis.

## Results

### Overview of CYTAP

CYTAP infers cytokine responses in individual cells from their transcriptional states. Because the same cytokine can induce distinct responses across cell types, CYTAP adapts pretrained cell-state embeddings to interpret responses in the context of the receiver cell type (Fig. 1 and Methods). Raw cell-state embeddings contain both baseline variation associated with cell type and donor, and response variation induced by cytokine stimulation (Fig. 1a). Subtracting the donor- and cell-type-matched control centroid yields a matched deviation that emphasizes response-associated variation but removes the corresponding cell-type baseline. To retain this context, CYTAP also constructs a donor-adjusted receiver state by subtracting the donor-specific control offset relative to the pooled control centroid of the same cell type. The two vectors are concatenated into a composite representation that combines receiver-cell context with the deviation from its matched baseline (Fig. 1b and Supplementary Note 1).

**Figure 1.**
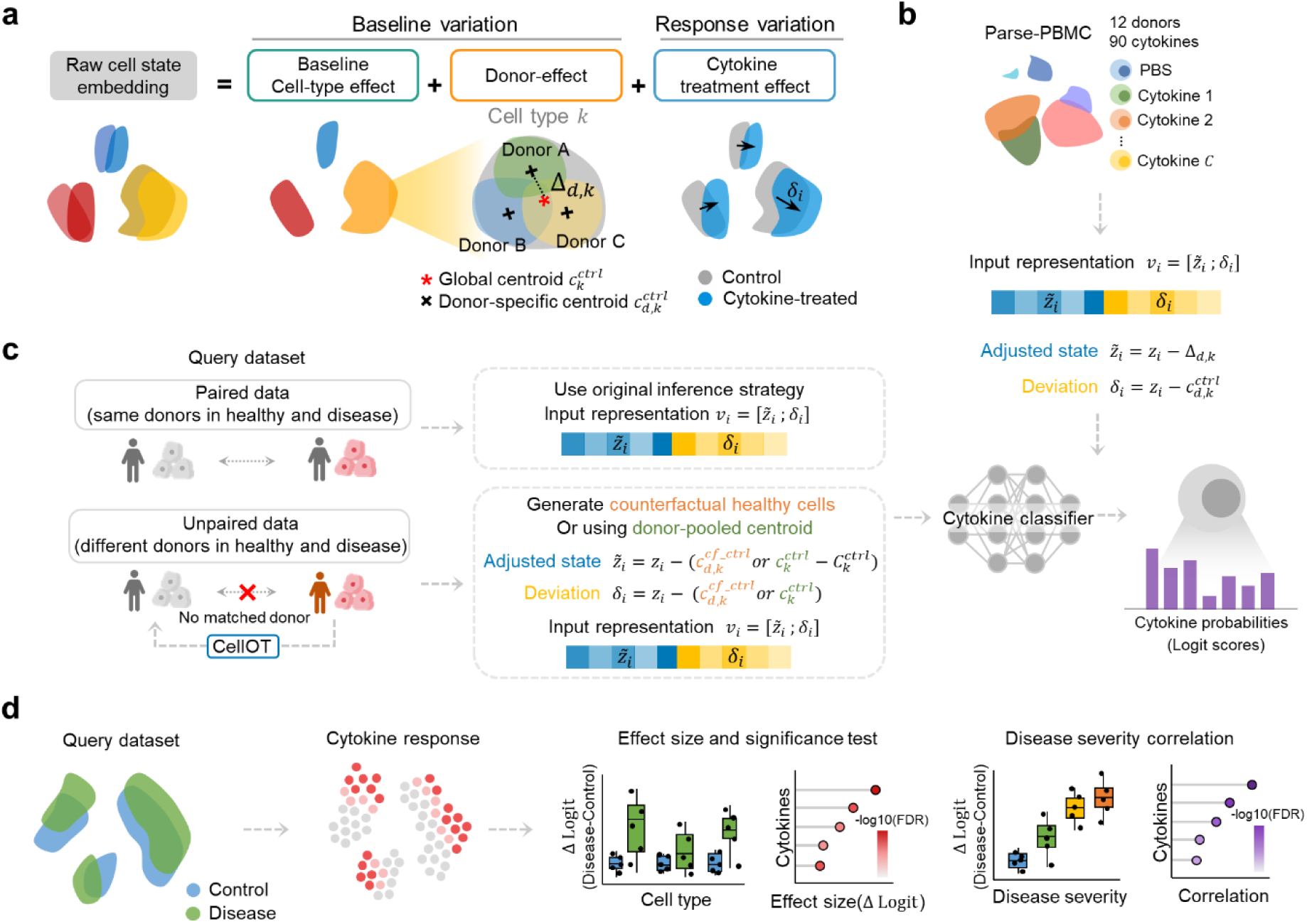
Overview of CYTAP. a, Structure of variation in cell-state embeddings. A raw embedding combines baseline variation, comprising a cell-type effect and a donor effect, with the response variation induced by cytokine treatment. Δ*_d_*_,*k*_, offset of the control centroid of donor 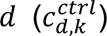 from the pooled control centroid of cell type 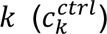; *δ_i_*, deviation of cell *i* from its donor- and cell-type-matched control centroid 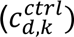. **b,** Model training. Each cell is represented by the composite representation *v_i_*, the concatenation of its adjusted state *~z_i_*, obtained by subtracting the estimated donor-associated baseline offset, and its deviation *δ_i_*. This representation was used to train a cytokine classifier on the Parse-PBMC dataset, which returns a probability for each of the 90 cytokines, reported as a logit score. **c,** Inference. For paired data, in which the same donors are sampled in the control and disease conditions, inputs are constructed as in training. For unpaired data, the donor-matched control centroid is replaced either by a counterfactual control centroid computed from control states generated with CellOT 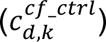 or by a donor-pooled control centroid of the same cell type 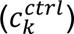. The resulting representation is passed through the same classifier. As control is not a class, probabilities are relative across the 90 cytokines, so condition effects are quantified as Δ logit relative to control. **d,** Downstream analysis. Per-cell logit scores are aggregated by donor and cell type to give cell-type-resolved cytokine responses. From these aggregated scores, the effect of a condition is quantified as an effect size (Δ logit, disease − control) and tested across donors. Cytokine responses can also be related to clinical variables, such as disease severity.

We trained a multiclass classifier on these composite representations using the Parse-PBMC cytokine perturbation dataset (Fig. 1b). For each cell, the classifier predicts probabilities across 90 cytokines, which are logit-transformed to obtain response scores. Control cells are used to define the baseline but are not included as an output class. The predicted probabilities therefore represent relative support among cytokine classes, and cytokine responses are identified as increases in response scores relative to control cells.

For query datasets with donor-matched controls, composite inputs are constructed as in training (Fig. 1c). When disease and control samples originate from different donors, CYTAP uses either a donor-pooled control centroid of the same cell type or a counterfactual control centroid computed for each donor and cell type from CellOT-generated control states^6^ (Supplementary Note 2). The counterfactual reference preserves the paired input formulation, whereas the pooled-reference input combines the raw receiver state with its deviation from the pooled control centroid (Methods). These alternatives allow the trained classifier to be applied to unpaired datasets.

Per-cell response scores can be visualized at single-cell resolution and summarized by donor and cell type (Fig. 1d). Condition effects are quantified as differences in the mean donor-level median scores between stimulated or disease samples and controls (Δ logit). The donor-level summaries can also be correlated with clinical variables, such as disease severity, to identify disease-associated cytokine-response patterns (Methods).

### Donor-adjusted receiver states and matched deviations improve cytokine-response prediction

To assess how each component of the composite representation contributes to cytokine-response prediction, we performed a donor-held-out input ablation using four-fold cross-validation on the Parse-PBMC dataset (Fig. 2a and Methods). Under the same training configuration, we trained classifiers on five inputs: log-normalized expression of 2,058 highly variable genes (HVGs), raw cell-state embeddings *z_i_*, donor-adjusted receiver states *~z_i_*, matched deviations *δ_i_*, and composite representations *v_i_*. At epoch 10, averaged across the four folds, all embedding-based inputs outperformed the HVG baseline in both macro F1 score (0.196–0.237 versus 0.096) and the area under the receiver operating characteristic curve (AUROC; 0.863– 0.884 versus 0.748). All three transformed inputs also outperformed the raw embedding (macro F1 score, 0.224–0.237 versus 0.196; AUROC, 0.877–0.884 versus 0.863). The composite representation performed similarly to the donor-adjusted state (macro F1 score, 0.237 versus 0.236; AUROC, 0.884 versus 0.883) and yielded slightly higher scores than the matched deviation (macro F1 score, 0.224; AUROC, 0.877; Fig. 2b,c). Thus, both components were predictive individually, and their combination retained performance comparable to that of the stronger component.

**Figure 2.**
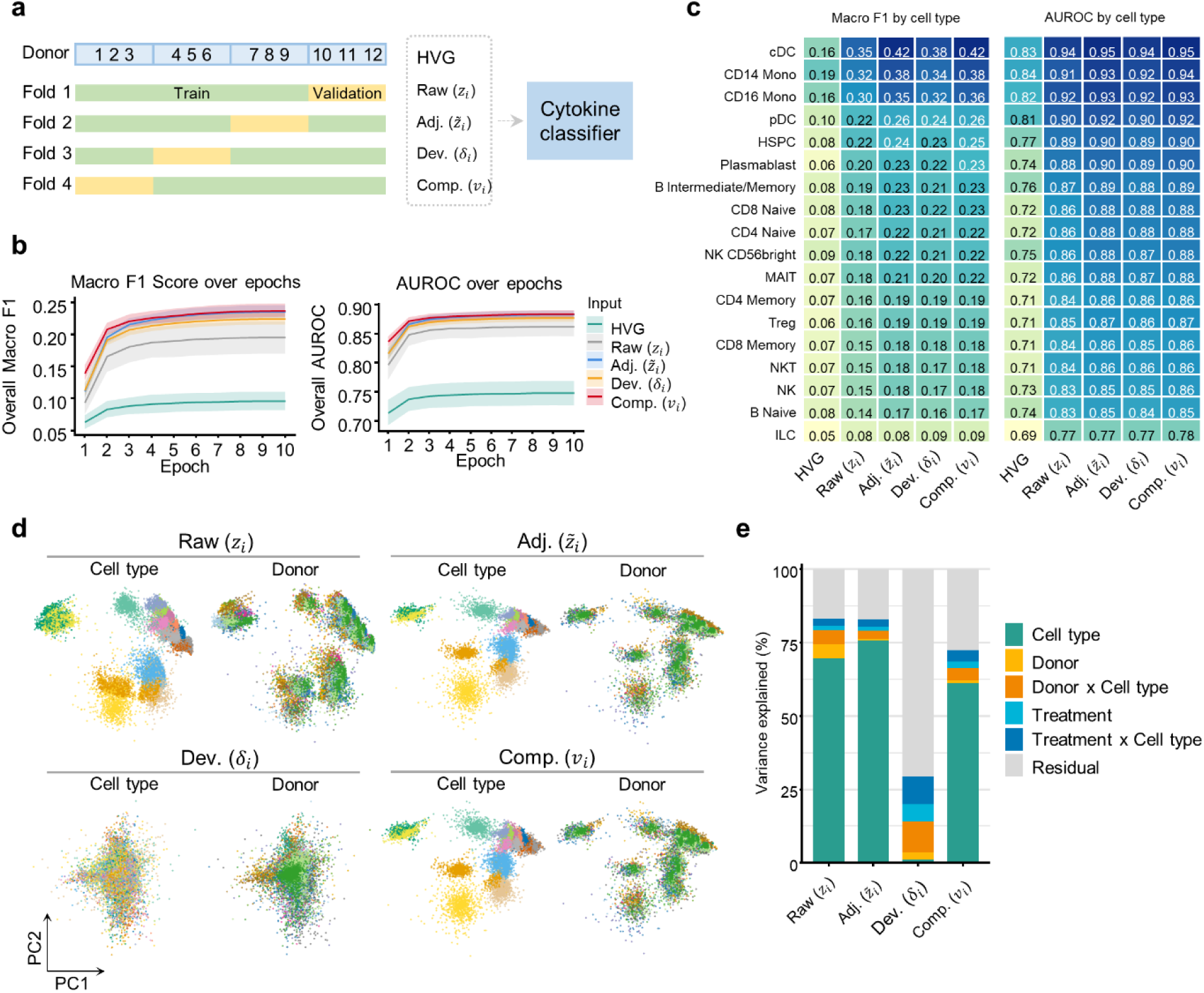
Input ablation for the cytokine classifier. a, Schematic of the experiment. The Parse-PBMC dataset was split into four folds by donor, so that the validation donors of each fold were unseen during training, and a cytokine classifier (MLP) was trained on each of five input representations: HVG (highly variable genes), Raw (*z_i_*, raw cell state), Adj. (*~z_i_*, donor-adjusted state), Dev. (*δ_i_*, matched deviation) and Comp. (*v_i_*, the composite representation). **b,** Macro F1 score and AUROC on the held-out validation donors over 10 training epochs. Lines, mean across the four folds. Shading, s.d. across the four folds. **c,** Heatmap of per-cell-type macro F1 and AUROC at epoch 10, averaged across the four folds. Cell types are ordered by their score under Comp. input representation. **d,** Centroids of each donor, cell type and cytokine in the Parse-PBMC data for Raw, Adj., Dev. and Comp., shown on the first two principal components. **e,** Hierarchical partitioning of variance for the four input representations. Bars show the contribution of cell type, donor, donor × cell type, treatment, treatment × cell type and residual variation, each expressed as a percentage of that representation’s own total variance. Donor × Cell type and Treatment × Cell type are interaction terms.

We next examined how these transformations redistributed variation in the embedding space. In principal component analysis (PCA) of donor–cell-type–cytokine centroids, the donor-adjusted state showed weaker donor-associated structure than the raw embedding, whereas the matched deviation showed no clear separation by cell type in the first two principal components (Fig. 2d). Hierarchical variance partitioning further distinguished the information retained by each representation (Fig. 2e and Methods). Relative to the raw embedding, the donor-adjusted state reduced the proportions of total variance attributable to donor and to donor-by-cell-type effects. The matched deviation instead increased the proportions attributable to treatment and to treatment-by-cell-type effects, while nearly eliminating the contribution of cell type. The composite representation retained cell-type structure alongside treatment-associated variation, providing receiver-cell context together with a response-enriched deviation.

To evaluate the dependence of prediction performance on the pretrained embedding, we compared raw embeddings from five single-cell foundation models using the same donor-held-out design (Extended Data Fig. 1a). State SE^1^ and scFoundation^3^ outperformed the HVG baseline in both metrics (macro F1 score, 0.196 and 0.146 versus 0.096; AUROC, 0.863 and 0.832 versus 0.748), with State SE achieving the highest scores. Geneformer v2^36^ matched the HVG baseline in macro F1 score (0.096) but exceeded it in AUROC (0.783), whereas scGPT^4^ and UCE^2^ scored below the baseline in both metrics (macro F1 score, 0.054 and 0.044; AUROC, 0.716 and 0.683; Extended Data Fig. 1b,c). These results supported the use of State SE as the embedding model for CYTAP under this evaluation setting.

We then tested whether the benefit of the input transformations extended to scFoundation, the other embedding model that outperformed the HVG baseline in both metrics. All three transformed inputs improved prediction relative to the raw scFoundation embedding (macro F1 score, 0.180–0.186 versus 0.146; AUROC, 0.854–0.857 versus 0.832). The composite representation again performed comparably to the better-performing individual component (macro F1 score, 0.186 versus 0.185 for the matched deviation and 0.180 for the donor-adjusted state; AUROC, 0.857 versus 0.855 and 0.854, respectively; Extended Data Fig. 1d,e). The relative performance of the two components differed between embedding models: the donor-adjusted state performed better than the matched deviation with State SE, whereas the two performed similarly with scFoundation. Across both embedding models, the composite representation maintained performance comparable to the better-performing component, supporting its use to combine receiver-cell context with matched perturbation deviations.

### CYTAP infers IFN-β responses despite sparse receptor transcript detection

We first applied CYTAP to PBMCs from eight patients with systemic lupus erythematosus (SLE), with cells from each donor stimulated in vitro with IFN-β for 6 h or left untreated^37^ (Fig. 3a). Transcripts of *IFNAR1* and *IFNAR2,* which encode the two subunits of the type I interferon receptor, were sparsely detected, whereas the interferon-stimulated genes *ISG15* and *IFIT1* were strongly induced upon treatment (Fig. 3b).

**Figure 3.**
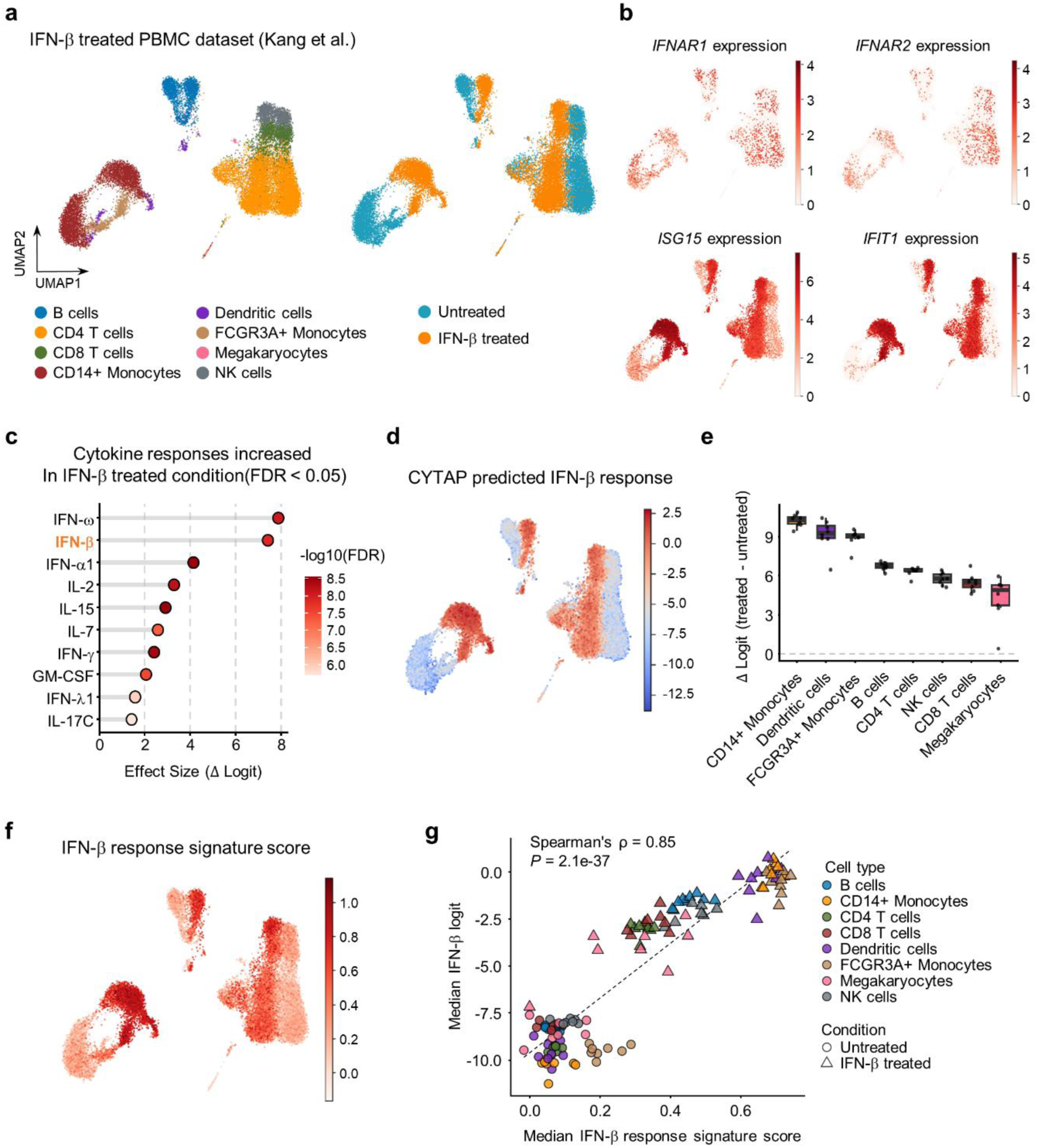
Validation in IFN-β-stimulated human PBMCs in vitro. a–g, PBMCs from eight patients with SLE, stimulated in vitro with IFN-β for 6 h, alongside an untreated control from each donor (paired). **a,** UMAP embedding colored by cell type and by stimulation. **b,** Expression of *IFNAR1* and *IFNAR2*, the IFN-β receptor subunits, and of the interferon-stimulated genes *ISG15* and *IFIT1* on the same embedding. **c,** Lollipop plot of the top 10 cytokines with significantly increased CYTAP-predicted response scores in IFN-β-treated versus untreated cells (FDR < 0.05; one-sided paired t-test with Benjamini–Hochberg correction, n = 8 donors), ranked by effect size (Δ logit), with IFN-β shown in bold orange. **d,** CYTAP-predicted IFN-β response on the UMAP embedding, colored from blue (low) to red (high logit score). **e,** Box plots of the CYTAP-predicted IFN-β response by cell type, expressed as the change in logit between each donor’s treated and untreated cells (Δ logit). Each dot is one donor (n = 8). Center line, median; box, interquartile range; whiskers, 1.5× interquartile range. **f,** IFN-β response signature score, calculated from the response to interferon-β gene set (GO:0035456), shown on the UMAP embedding. Darker red indicates a higher score. **g,** Median signature score plotted against the median CYTAP IFN-β logit score, calculated for each donor, cell type and condition. Spearman’s correlation coefficient (ρ) and the corresponding *P* value are shown.

In donor-level comparisons of treated and untreated samples, CYTAP ranked IFN-β second by effect size (Δ logit) among cytokines with significantly increased response scores (FDR < 0.05), after IFN-ω, another type I interferon that signals through the same receptor complex (Fig. 3c). IFN-β response scores were elevated in treated cells at single-cell resolution (Fig. 3d). The magnitude of this increase varied across cell types, with CD14+ monocytes showing the largest median donor-level change (Fig. 3e).

To assess concordance with the well-characterized IFN-β transcriptional response^38^, we scored each cell using the Gene Ontology “response to interferon-β” gene set (GO:0035456; Methods). The signature score increased after treatment (Fig. 3f) and was strongly correlated with the CYTAP IFN-β response score when both measures were summarized as medians for each donor-cell-type-condition group (Spearman’s ρ = 0.85, *P* = 2.1 × 10^-37^; Fig. 3g). Together, these results show that CYTAP prioritizes the applied cytokine and captures cell-type variation in the associated response despite sparse detection of receptor transcripts.

### CYTAP distinguishes cytokine responses in stimulated macrophages

We next applied CYTAP to PBMC-derived macrophages from four donors treated in vitro with IFN-β, IFN-γ, IL-4, TNF-α or TNF-α plus IFN-γ, alongside donor-matched untreated controls^39^ (Fig. 4a). Predicted response scores for IFN-β, IFN-γ and IL-4 were highest under the corresponding stimulation conditions, with elevated IFN-γ response scores following treatment with IFN-γ either alone or in combination with TNF-α (Fig. 4b–d). Among cytokines with significantly increased response scores (FDR < 0.05), the applied cytokine ranked first by effect size for IFN-γ and IL-4 and second for IFN-β, after IFN-α1, which signals through the same receptor complex (Fig. 4e–g). In IL-4-treated cells, only the IL-4 and IL-13 response scores increased significantly (Fig. 4g), consistent with the shared use of the type II receptor complex IL-4Rα-IL-13Rα1 by these cytokines^40^. By contrast, no cytokine response score increased significantly in TNF-α-treated cells relative to donor-matched untreated controls. Although the TNF-α response score was higher in this condition than in the other cytokine-treated conditions, its control-relative effect size remained close to zero (Extended Data Fig. 2b). TNF-α-treated cells also overlapped untreated cells in both the cell-state and HVG-based embeddings and showed no significant increase in the TNF-α signature score (Fig. 4a and Extended Data Fig. 2a,c).

**Figure 4.**
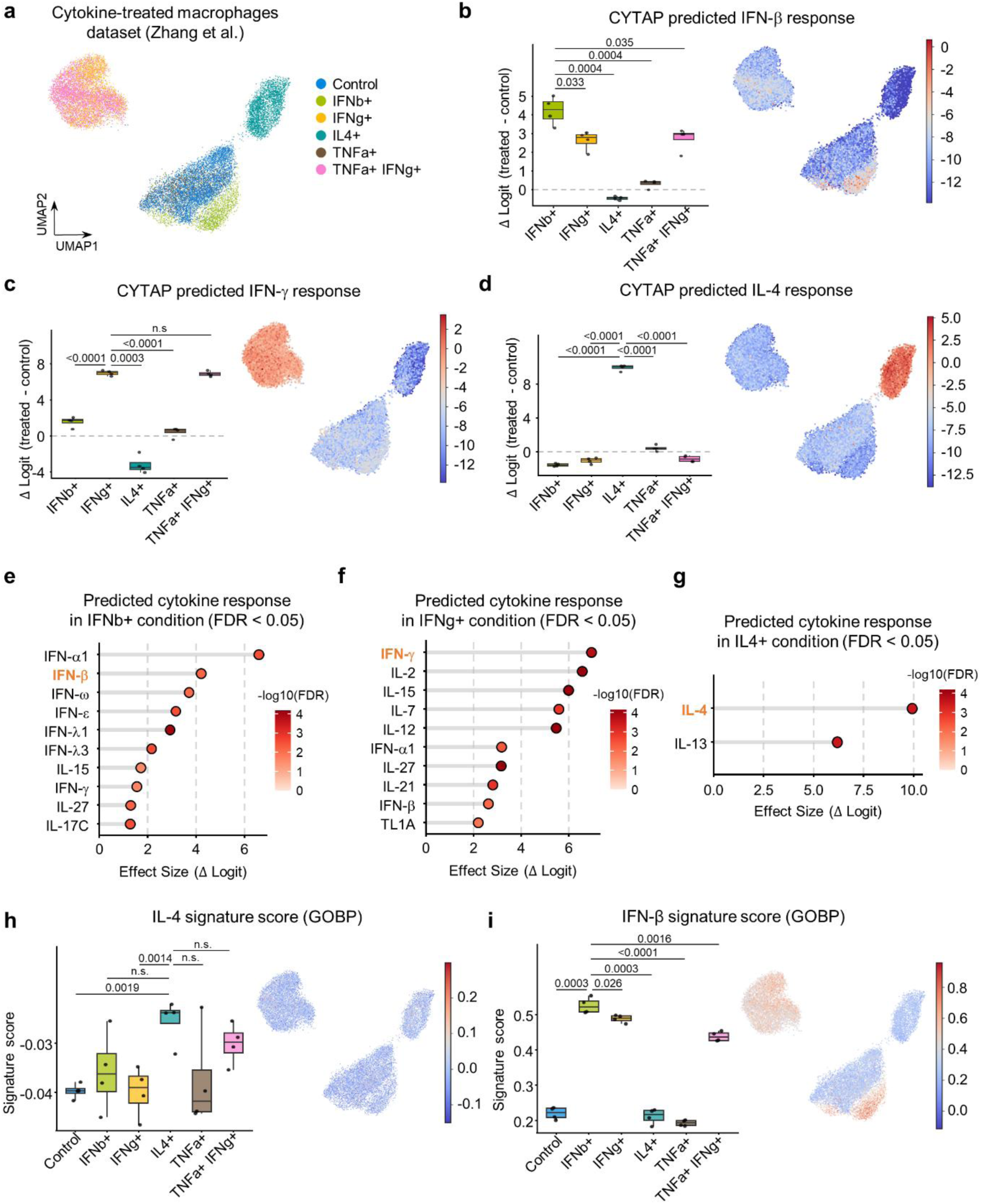
Validation in cytokine-treated human macrophages in vitro. a–i, Macrophages differentiated from the PBMCs of four donors and treated in vitro with IFN-β, IFN-γ, IL-4, TNF-α or TNF-α plus IFN-γ, alongside an untreated control (paired; n = 4 donors). Center line, median; box, interquartile range; whiskers, 1.5× interquartile range. **a,** UMAP embedding colored by condition. **b–d,** CYTAP-predicted response to IFN-β (b), IFN-γ (c) and IL-4 (d). Box plots (left) show the response in each treatment condition, expressed as the change in logit relative to each donor’s control (Δ logit). Each dot is one donor. UMAP embeddings (right) are colored by the predicted logit score. *P* values are from one-sided paired t-tests across four donors (n.s., *P* > 0.05). **e–g,** Lollipop plots of up to 10 cytokines with significantly increased CYTAP-predicted response scores in cells treated with IFN-β (e), IFN-γ (f) or IL-4 (g) versus control (FDR < 0.05; one-sided paired t-test with Benjamini–Hochberg correction, n = 4 donors), ranked by effect size (Δ logit), with the treated cytokine shown in bold orange. **h,i,** Response signature scores by condition for IL-4 (h; response to interleukin-4, GO:0070670) and IFN-β (i; response to interferon-β, GO:0035456), shown as box plots (left) and on UMAP embeddings (right). Darker red indicates a higher score. *P* values are from one-sided paired t-tests across four donors (n.s., *P* > 0.05).

We then compared CYTAP predictions with cytokine response signature scores derived from curated gene sets (Methods). The IL-4 signature score increased significantly in IL-4-treated cells relative to untreated controls and IFN-γ-treated cells but did not differ significantly from the scores in IFN-β, TNF-α or TNF-α plus IFN-γ treated cells (Fig. 4h). In contrast, CYTAP IL-4 response scores were higher in IL-4-treated cells than in each of the four other treatment conditions (all *P* < 0.0001; Fig. 4d), indicating a more selective treatment-associated response pattern in this dataset. For the interferons, both CYTAP and gene signature scores were highest under the corresponding stimulation conditions, although overlapping responses were observed across interferon treatments (Fig. 4b,c,i and Extended Data Fig. 2d). Thus, the signature scores also distinguished the two interferon treatment conditions, whereas the clearest additional discrimination provided by CYTAP was observed for IL-4.

Finally, we examined the contribution of the composite representation in this external dataset by applying classifiers trained with each of the four alternative inputs (Extended Data Fig. 2e–h). Among the alternative inputs, only the matched deviation placed IFN-β within the top ten cytokines in IFN-β-treated cells, ranking it third after the type I interferons IFN-α1 and IFN-ω. With the raw embedding, the IFN-β response score was highest in IFN-β-treated cells when compared across treatment conditions, but IFN-β did not rank among the top ten cytokines within that condition. With the HVG baseline and the donor-adjusted state, IFN-β response scores were higher in other conditions than in IFN-β-treated cells. The composite representation therefore ranked the applied IFN-β higher than any alternative input in this external macrophage dataset, supporting the combination of receiver-cell context and matched deviations for response inference beyond the training dataset.

### CYTAP recovers administered cytokines in vivo

To evaluate whether CYTAP could recover cytokine responses following in vivo administration, where cells may also be exposed to secondary signals from surrounding cells, we analyzed two paired datasets sampled before and after treatment. These datasets comprised bone marrow hematopoietic cells and peripheral blood T cells, allowing us to assess transfer to in vivo settings and, in the former case, beyond the PBMC training context.

The G-CSF dataset comprised bone marrow hematopoietic cells from two healthy donors sampled before and after G-CSF treatment^41^ (Fig. 5a). Transcripts of *CSF3R*, which encodes the G-CSF receptor, were detected predominantly in the myeloid lineage (Fig. 5b). Given the limited number of donors, we ranked cytokines by effect size without significance filtering. CYTAP ranked G-CSF ninth among all 90 cytokines in myeloid cells, and the predicted G-CSF response scores were highest in this lineage, consistent with the *CSF3R* expression pattern (Fig. 5c).

**Figure 5.**
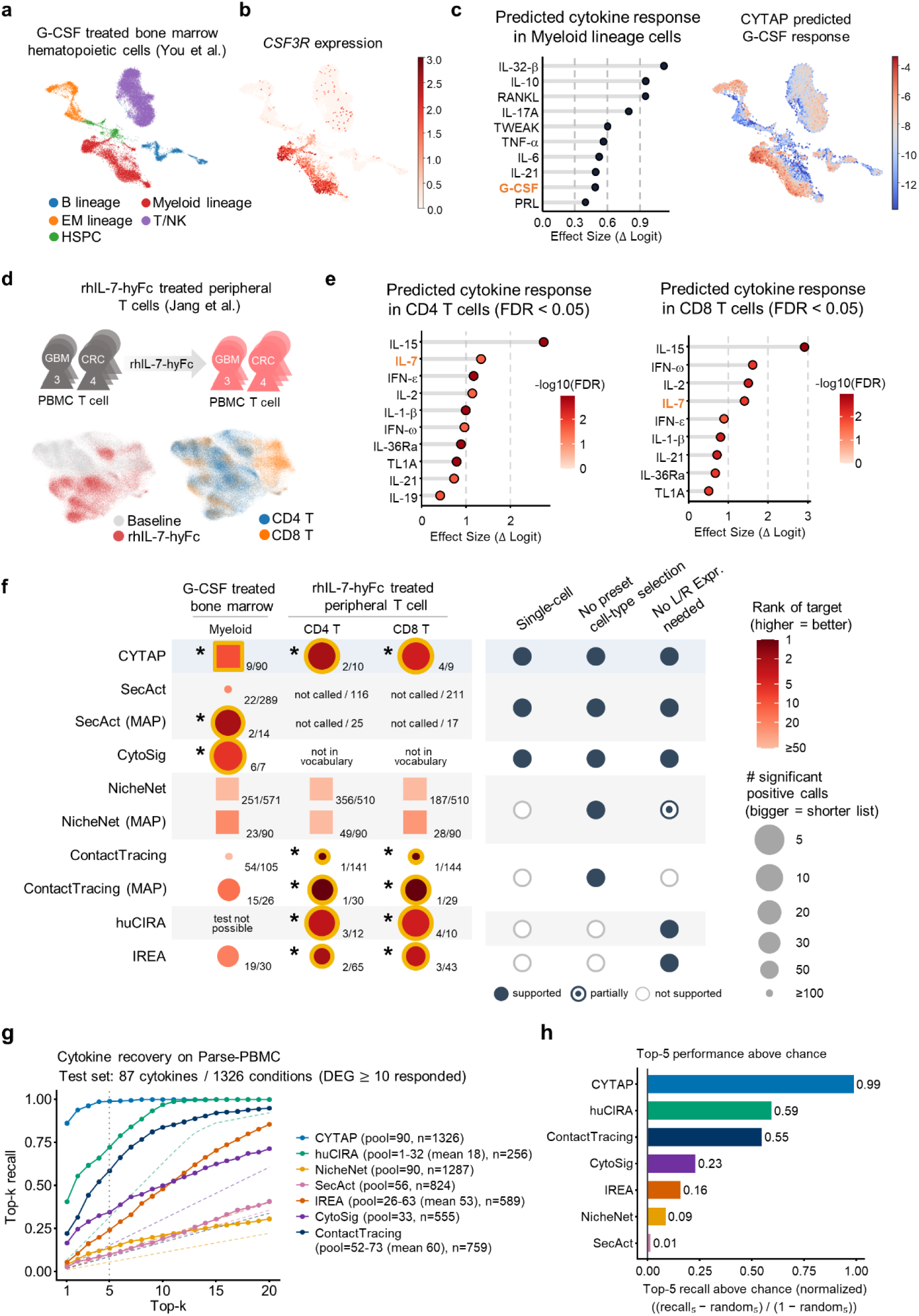
In vivo validation of CYTAP and benchmarking against existing cytokine-inference methods. a–c, Bone marrow cells from two healthy donors before and after G-CSF treatment (paired). **a,** UMAP embedding colored by cell type. **b,** Expression of *CSF3R*, the G- CSF receptor. **c,** Top 10 cytokines with the largest increase in CYTAP-predicted response (Δ logit) in myeloid lineage cells after G-CSF treatment (left; G-CSF in orange; ranked without significance testing, n = 2 donors), and the predicted G-CSF response on the UMAP embedding (right). **d,e,** Peripheral T cells from patients with glioblastoma (GBM; n = 3) or colorectal cancer (CRC; n = 4) before and after rhIL-7-hyFc treatment. **d,** Study design (top) and UMAP embedding colored by condition or by cell type (bottom). **e,** Cytokines with increased CYTAP-predicted response (Δ logit) after treatment (FDR < 0.05; one-sided paired t-test with Benjamini–Hochberg correction, n = 7 patients) in CD4 (left) and CD8 (right) T cells, with IL-7 in orange. **f,** Rank of the administered cytokine assigned by each cytokine-inference method in the G-CSF (**a–c**) and rhIL-7-hyFc (**d,e**) datasets. MAP, candidates restricted to ligands or receptors of the 90 CYTAP cytokines. Labels indicate the target rank and the number of significant positive calls; color, target rank (redder, better); circle size, inversely scaled to the number of significant calls; squares, rank without significance filtering; yellow ring with asterisk, target within the top 10. Scoring details and the right-hand criteria are defined in Methods. **g,** Top-k recall (k = 1–20) of the administered cytokine across cell type–cytokine conditions in the Parse-PBMC dataset. Dashed lines, random expectation for each method; pool, number of ranked candidates (range and mean where variable); n, number of evaluable conditions; dotted line, k = 5. **h,** Top-5 recall above chance (0, random; 1, perfect recovery).

We next analyzed peripheral blood T cells from seven patients, three with glioblastoma and four with colorectal cancer, sampled before and after treatment with rhIL-7-hyFc, a long-acting form of IL-7^42^ (Fig. 5d). Among cytokines with significantly increased response scores after treatment (FDR < 0.05), IL-7 ranked second of 10 in CD4+ T cells, after IL-15, and fourth of nine in CD8+ T cells, after IL-15, IFN-ω and IL-2 (Fig. 5e). The prioritization of IL-15 and IL-2 alongside IL-7 is consistent with their shared use of the common γ-chain receptor subunit and overlapping T-cell response programs^43,44^. Together, these results show that CYTAP prioritizes administered cytokines in the expected responding cell populations across both in vivo datasets.

### CYTAP recovers administered cytokines more consistently than existing methods

We compared CYTAP with six existing cytokine-inference methods, SecAct, CytoSig, NicheNet, ContactTracing, huCIRA and IREA, in the two in vivo datasets, using the rank assigned to the administered cytokine as the primary evaluation metric (Fig. 5f and Extended Data Figs. 3 and 4). Each method was evaluated using its respective candidate vocabularies and applicable significance filters, with additional comparisons restricting candidates to ligands or receptors corresponding to the 90 CYTAP cytokines (Methods). In the G-CSF dataset, with myeloid cells as the receiver population, SecAct ranked G-CSF second among 14 significant calls after candidate restriction, and CytoSig ranked it sixth among seven. NicheNet, ContactTracing and IREA did not place G-CSF within their top ten candidates, whereas huCIRA could not be applied because too few genes overlapped its reference dictionary (Extended Data Fig. 3a– e).

In the IL-7 dataset, ContactTracing, huCIRA and IREA placed IL-7 within the top ten in both CD4+ and CD8+ T cells. For CD4+ and CD8+ T cells, respectively, the ranks and numbers of significant calls were 1/30 and 1/29 for ContactTracing after candidate restriction, 3/12 and 4/10 for huCIRA, and 2/65 and 3/43 for IREA. SecAct inferred decreased IL-7 activity after treatment and therefore did not include IL-7 among its positive calls. NicheNet, which ranks ligands without significance testing, did not place IL-7 within the top ten, whereas CytoSig did not include IL-7 in its candidate vocabulary (Extended Data Fig. 4a–e).

Under these evaluation settings, CYTAP was the only method evaluated that placed the administered cytokine within the top ten in both in vivo datasets (Fig. 5c,e,f). Its G-CSF ranking was based on effect size without significance filtering, given the limited number of donors. In the IL-7 dataset, CYTAP returned ten significant calls in CD4+ T cells and nine in CD8+ T cells, fewer than the other methods that placed IL-7 within the top ten. CYTAP also generates single-cell scores across annotated cell types using a shared classifier, without requiring a receiver population to be selected in advance or filtering candidate cytokines by ligand or receptor expression.

We evaluated Scupa separately because it quantifies predefined polarization states rather than individual cytokine responses. In the G-CSF dataset, three of the five monocyte states showed higher polarization scores after treatment (Extended Data Fig. 3f), although none of these states was annotated as G-CSF-induced in the reference. In the IL-7 dataset, the largest increase among CD4+ T-cell states occurred in T4.f, one of the states induced by IL-7, whereas most CD8+ T-cell states increased after treatment (Extended Data Fig. 4f). Thus, Scupa captured treatment-associated polarization changes, but its state-level outputs did not directly rank the administered cytokine.

To extend the comparison across a broader range of cytokines, we evaluated recovery of the applied cytokine in 1,326 responsive cell type–cytokine conditions spanning 87 cytokines in the Parse-PBMC dataset. Responsive conditions were defined as combinations with at least 10 differentially expressed genes relative to PBS-treated cells (Methods). CYTAP predictions were obtained from the cross-validation fold in which each cell’s donor was held out, and each method was evaluated only on conditions for which the applied cytokine was represented in its candidate pool. CYTAP achieved the highest top-k recall across the evaluated range of k and recovered the applied cytokine within the top five in nearly all evaluable conditions (Fig. 5g). To account for differences in candidate-pool size, we also calculated top-5 recall normalized relative to each method’s chance expectation. CYTAP ranked highest by this measure, followed by huCIRA and ContactTracing (Fig. 5h). Although the donors used for CYTAP evaluation were held out from classifier training, CYTAP was trained on the same perturbation resource, and the huCIRA reference was derived from that resource. This benchmark therefore assesses performance within the Parse-PBMC distribution and complements the independent in vivo comparisons above.

### CYTAP identifies severity-associated cytokine responses in COVID-19

When donor-matched controls are unavailable, CYTAP requires an alternative reference to construct its input representation. We evaluated cell-type-matched pooled control centroids and CellOT-derived counterfactual control centroids in held-out donors from the Parse-PBMC cross-validation, using experimentally observed donor- and cell-type-matched PBS centroids as the benchmark reference. To assess whether the relative performance of pooled and counterfactual references depended on the control population used, we constructed each reference using PBS-treated cells from either the held-out donors or the training donors. The fold-specific cytokine classifiers were kept fixed. In both CellOT settings, the source population comprised cytokine-treated cells from the held-out donors, whereas the target PBS population differed. Transported states were averaged within each held-out donor and cell type to obtain counterfactual control centroids (Methods and Supplementary Note 2). With held-out-donor controls, the counterfactual reference approached matched-control performance (overall AUROC, 0.88 and 0.88; macro F1 score, 0.23 versus 0.24), whereas the pooled reference yielded lower scores (AUROC, 0.87; macro F1 score, 0.21; Extended Data Fig. 5a). Using training-donor controls reduced overall performance for both alternatives, but the counterfactual reference retained higher overall scores than the pooled reference (AUROC, 0.87 versus 0.86; macro F1 score, 0.21, versus 0.19). Thus, counterfactual references improved overall prediction relative to pooling under both control-population settings, although absolute performance depended on the control population and the benefit varied across cell types.

In a separate variance-partitioning analysis using all 12 donors, donor and donor-by-cell-type effects accounted for larger proportions of total variance with pooled than with matched references, particularly in the deviation component. Counterfactual references showed smaller relative contributions of these donor-related effects than pooled references (Extended Data Fig. 5b). These results supported the use of counterfactual rather than pooled control references for subsequent unpaired analyses. For each query dataset, we therefore fitted CellOT maps from disease cells to the observed control population and used the resulting donor- and cell-type-specific counterfactual centroids to construct the classifier inputs.

We applied CYTAP to PBMCs from 73 donors in the COVID-19 Multi-omic Blood Atlas (COMBAT) cohort^45^, comprising healthy volunteers and patients with mild, severe or critical COVID-19 without donor-matched controls (Fig. 6a). Among cytokines with response scores positively associated with severity (FDR < 0.05), IL-32β and IL-21 ranked first and second by the Spearman correlation between donor-level median scores and ordinal severity (Fig. 6b). IL-32β response scores increased with severity in most cell types (Fig. 6c), despite the lack of a defined specific receptor for IL-32^46,47^. Previous studies have reported elevated serum IL-21 in severe COVID-19 and persistently elevated circulating IL-32 after infection^48–51^. Other top-ranked cytokines, including IL-6, IL-1β and IL-8, have also been associated with severe COVID-19^52,53^.

**Figure 6.**
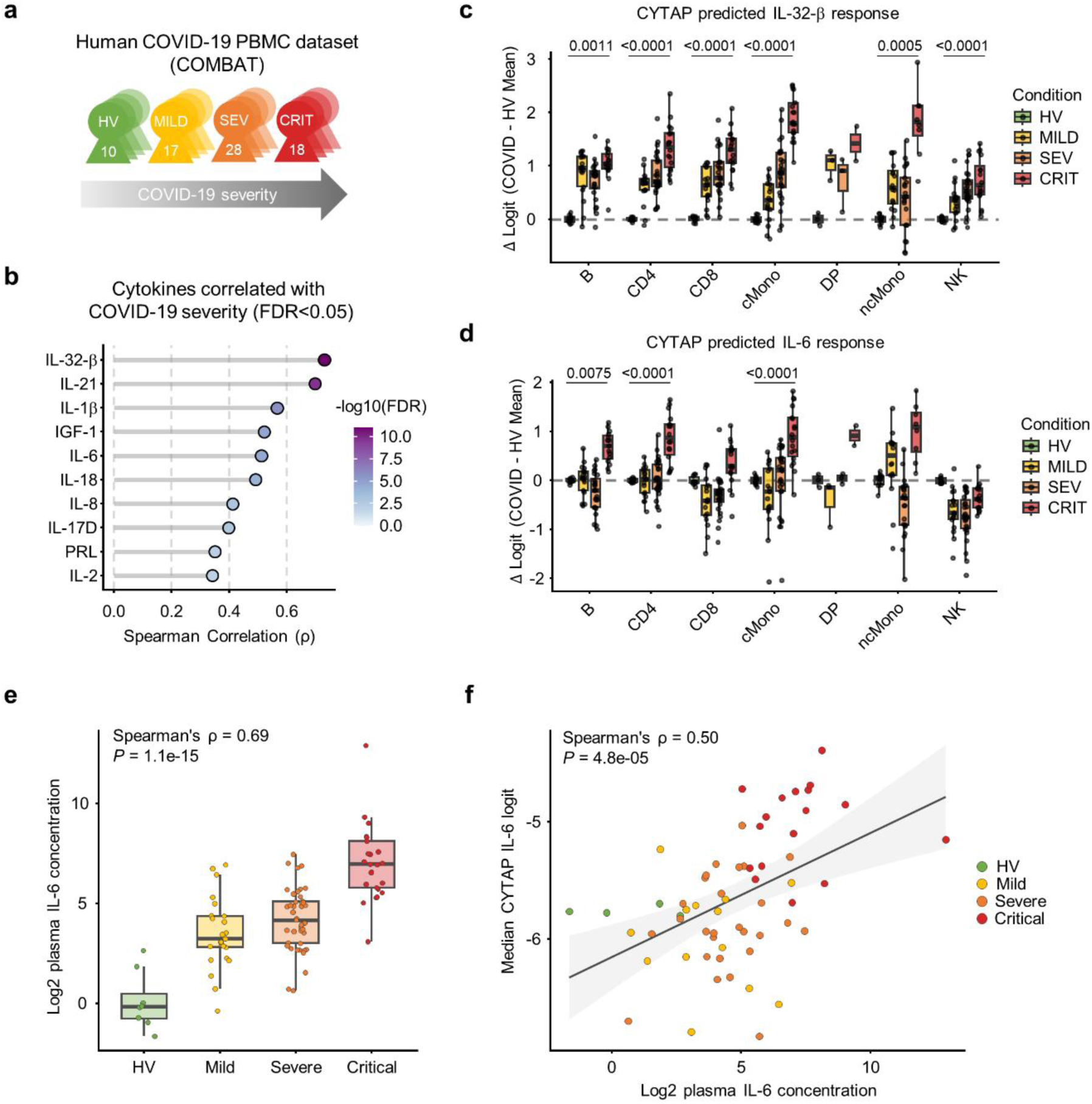
Application to an unpaired COVID-19 cohort. a, Study design. PBMCs from 73 unpaired donors in the COVID-19 Multi-omic Blood Atlas (COMBAT) cohort across four groups: healthy volunteers (HV; n = 10), mild (MILD; n = 17), severe (SEV; n = 28) and critical (CRIT; n = 18) COVID-19. **b,** Top 10 cytokines whose CYTAP-predicted response increases with COVID-19 severity (Benjamini–Hochberg FDR < 0.05 for the Spearman correlation with ordinal severity, with a positive regression slope; n = 73 donors), ranked by Spearman’s ρ. Correlations used the median logit per donor. **c,d,** Box plots of the CYTAP-predicted response to IL-32β (c) and IL-6 (d) by cell type. Δ logit, median logit of each donor relative to the mean of HV donors (HV shown for reference). Each dot is one donor. FDR (Benjamini–Hochberg-adjusted Spearman P value) is shown for cell types with a significant correlation with severity. Center line, median; box, interquartile range; whiskers, 1.5× interquartile range. **e,** Plasma IL-6 concentration (Luminex) by severity group in all COMBAT donors with a plasma sample, including those without scRNA-seq data. Each dot is one donor. Spearman’s ρ and *P* value were calculated against the ordinal severity score. Donors whose plasma IL-6 fell below the lower limit of detection (17 of 25 healthy volunteers and 5 of 98 patients) were excluded from the plot and the correlation analysis (n = 101 donors). **f,** Median CYTAP-predicted IL-6 response per donor versus plasma IL-6 concentration, colored by severity group, with Spearman’s ρ and *P* value. Donors below the lower limit of detection were excluded, as in **e** (n = 61 donors).

To compare inferred responses with independently measured cytokine protein concentrations, we analyzed the COMBAT plasma Luminex panel. Because the panel did not include IL-21 or IL-32, we focused on IL-6. The predicted IL-6 response scores increased significantly with severity in B cells, CD4+ T cells and classical monocytes (Fig. 6d). Plasma IL-6 concentrations also increased with severity (Fig. 6e). Among donors with both scRNA-seq data and quantifiable plasma IL-6 measurements, the donor-level median CYTAP IL-6 response score correlated positively with plasma IL-6 concentration (n = 61; Spearman’s ρ = 0.50, *P* = 4.8 × 10^-5^; Fig. 6f). Repeating inference with the pooled control reference also prioritized IL-32β and IL-21, and IL-6 remained among the top ten cytokines, showing that these severity associations were retained without CellOT (Extended Data Fig. 6).

Receptor transcript abundance showed a different pattern. Despite the increases in predicted IL-6 response and plasma IL-6 concentration, *IL6R* expression tended to decline with severity, most notably in classical monocytes, which had the highest *IL6R* expression among the cell types examined (Extended Data Fig. 7a). *IL6* transcripts were rarely detected in most cell types. The IL-6/JAK/STAT3 signature score was elevated mainly in classical and non-classical monocytes (Extended Data Fig. 7b). Together, these results link CYTAP-inferred IL-6 responses to disease severity and circulating IL-6 protein concentrations without a corresponding increase in receptor transcript abundance.

## Discussion

In this study, we present CYTAP, a framework for inferring cytokine responses from the transcriptional states of receiver cells. By combining a donor-adjusted receiver state with its deviation from a matched baseline, CYTAP learns cell-type-dependent cytokine-response patterns in a pretrained embedding space and transfers them across donors and datasets. It assigns response scores to individual cells without requiring detection of ligand or receptor transcripts. Recovery of the administered G-CSF and IL-7 responses further indicates that the cytokine source need not be represented by a sender-cell population in the profiled samples. Given the accessibility of blood-derived immune cells in clinical studies^54,55^, this raises the possibility of using blood transcriptomes to identify cytokine-associated responses to signals originating outside the sampled compartment and to prioritize candidate signaling pathways for functional investigation.

In vitro, CYTAP recovered the IFN-β response in SLE PBMCs despite sparse receptor detection and a shorter stimulation period than in the training data (6 h versus 24 h in healthy PBMCs). In macrophages, CYTAP detected the IL-4 response and showed higher IL-4 scores in the IL-4-treated condition than in each of the other cytokine-treated conditions examined. The IL-4 gene signature also increased relative to untreated controls, but did not significantly distinguish IL-4 treatment from several alternative cytokine treatments. Both CYTAP and gene signature scores distinguished the corresponding interferon treatment conditions, although overlapping responses remained apparent. The representation used for inference was also important. Baseline-relative transformations improved held-out-donor prediction relative to raw embedding and HVGs, with the composite representation performing comparably to the better-performing individual component in both State SE and scFoundation. In the external macrophage dataset, the composite input ranked the applied IFN-β higher than either component alone. Together, these findings support combining receiver-cell context with matched deviations as a strategy for transferring cytokine-response inference across datasets.

The in vivo benchmarks also highlighted differences in the information used by existing methods. ContactTracing, which compares receptor-expressing and receptor-null cells, ranked the IL-7-associated receptor signal first, whereas SecAct inferred a negative change in IL-7 activity. These contrasting outputs illustrate that receptor-associated and reference-signature approaches can yield different interpretations of the same perturbation. IL-7-mediated suppression of *IL7R* transcription^56^ and negative feedback in cytokine signaling^57^ indicate that cytokine exposure can reduce the expression of feedback-regulated genes, which could lower activity estimates from reference signatures that include them. However, the present analyses do not establish whether such feedback contributed to the negative SecAct prediction. CYTAP instead prioritizes cytokines through similarity to experimentally stimulated cells, without requiring ligand or receptor detection, and therefore provides a complementary readout.

The ability to summarize single-cell response scores by donor also allows CYTAP to examine clinical severity gradients without limiting the analysis to a single case-control comparison. In the unpaired COMBAT cohort, CYTAP identified severity-associated IL-32β and IL-21 responses, neither of which could be assessed using the available Luminex panel. IL-32β was prioritized despite lacking a well-defined receptor^46,47^, a case that receptor-based approaches cannot readily address. For IL-6, the predicted response was positively associated with independently measured plasma protein concentrations among donors with quantifiable measurements, providing evidence of concordance across measurement modalities. This association was observed despite sparse *IL6* transcript detection and a tendency for *IL6R* expression to decline with severity, indicating that the predicted IL-6 response did not track ligand or receptor transcript abundance. This discordance may reflect IL-6 trans-signaling^27^, although this mechanism was not tested here. More broadly, these clinical associations help interpret disease-associated cell states in terms of cytokine response programs but do not identify causal cytokine drivers.

Several limitations should be considered. First, predictions depend on the pretrained embedding, the cell-type annotations used to construct reference centroids and the control reference available for each dataset. CellOT-derived centroids are model-imputed references rather than observed donor-matched controls and do not by themselves eliminate confounding in unpaired comparisons. The recovery of the same leading cytokines using a pooled reference in COMBAT supports the robustness of those leading associations, but does not establish that all predictions are independent of reference choice. Second, the current response dictionary is restricted to the 90 cytokines profiled in the human PBMC training atlas. Validation in blood-derived macrophages and bone marrow extends the evaluation to additional human immune contexts, but applicability to non-immune cells and other species remains untested. Reported differences between human and mouse cytokine responses^32^ further motivate dedicated validation before cross-species application. Third, cytokines with overlapping response programs may not be uniquely distinguishable, particularly when they share receptor complexes, as with type I interferons or IL-4 and IL-13. Responses to perturbations outside the dictionary may also be assigned to profiled cytokines with similar programs. Because control cells define the baseline but are not included as an output class, the predicted probabilities represent relative support among cytokine classes rather than the probability of a response versus no response. Condition-associated score changes should therefore be interpreted as evidence consistent with cytokine responses, not as proof of exposure to a particular cytokine. Expanding perturbation atlases across cytokines, cell types and species may address some of these coverage limitations.

These findings have broader implications for the use of single-cell foundation models in functional inference. Large-scale perturbation atlases and pretrained models are expanding rapidly^1–5,58–60^, yet benchmarks have shown that zero-shot embeddings do not consistently outperform simpler representations^61^ and that deep learning models do not necessarily outperform linear baselines in perturbation-effect prediction^62^. Our results address the task-aligned use of such representations: defining perturbation-associated deviations relative to control baselines improved cytokine-response prediction while keeping the pretrained encoder fixed. A shared embedding model provides a common feature space across datasets, although this alone does not guarantee removal of dataset-specific variation. The improvements observed with both State SE and scFoundation suggest that the input transformations are not restricted to a single encoder, while their effectiveness remains dependent on the biological information retained in the embedding. As perturbation references and representations improve, receiver-response approaches such as CYTAP could complement ligand-receptor-based communication analysis by providing evidence of cytokine-associated cell-state changes, particularly in clinical cohorts.

## Methods

We developed CYTAP to infer cytokine responses in individual cells by assessing whether receiver-cell state changes are consistent with experimentally induced cytokine responses. Each cell was represented by an embedding from the pretrained State embedding (SE) model. A cytokine-response classifier was trained on the Parse-PBMC dataset using a composite representation that concatenates the donor-adjusted receiver state with the cell’s deviation from its donor- and cell-type-matched PBS control centroid. The trained classifier was applied to query datasets to obtain per-cell cytokine response probabilities. These probabilities were logit-transformed and summarized as median scores for each donor, cell type and condition. The resulting donor-level summaries were used to quantify differences between conditions and assess association with disease severity.

### Generation of cell embeddings and model input

Each cell *i* was represented by an embedding vector *z_i_* ∈ ℝ*^D^*, where *D* = 2058, obtained from the pretrained State embedding (SE) model^1^. Gene expression counts were normalized to a total of 10,000 per cell and log-transformed. Embeddings were extracted using the SE-600M checkpoint and the “state emb transform” command of the official implementation. CYTAP was trained on the Parse-PBMC dataset, which contains PBS-treated control and cytokine-treated cells from the same donors. For each donor *d* and annotated cell type *k*, the donor- and cell-type-matched control centroid was defined as

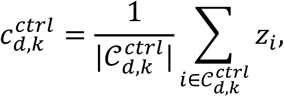

where 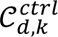 denotes the set of PBS control cells from donor *d* and cell type *k*. The pooled control centroid for cell type *k* was computed as

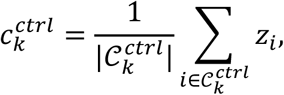

where 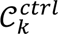 denotes the set of PBS control cells of cell type *k* pooled across the training donors. The donor-associated baseline offset relative to the pooled control reference was estimated as 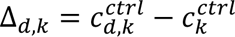 For each cell *i*, with donor *d*(*i*) and annotated cell type *k*(*i*), we constructed a donor-adjusted receiver state,

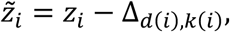

and a matched deviation,

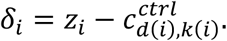

The donor-adjusted state subtracts the estimated donor-associated baseline offset, whereas the matched deviation represents the cell’s departure from its donor- and cell-type-matched control baseline. The final classifier input was the concatenation of these two vectors:

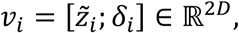

where the semicolon denotes vector concatenation. The rationale for this representation, based on a decomposition of each embedding into cell-type, donor and cytokine-response components, is given in Supplementary Note 1.

### Model training and inference

A multilayer perceptron (MLP) classifier was trained on the composite input vectors *v_i_* of cytokine-treated cells in the Parse-PBMC dataset. The network comprised two hidden layers with 2,048 and 1,024 units, respectively, each using batch normalization, ReLU activation and dropout at a rate of 0.3. The output layer contained 90 units, one for each cytokine. A softmax function converted the output logits into predicted probabilities *p_i_*_,*c*_ for cytokine *c* ∈ T, where T denotes the set of cytokine classes. PBS control cells were used to construct control reference centroids but were not included as a classifier output class. The predicted probabilities therefore represent relative support among the 90 cytokine classes rather than probabilities of response versus no response.

To emphasize difficult-to-classify cells, the classifier was optimized using multi-class focal loss with a focusing parameter of *γ*_focal_ = 2^63^. Training used the AdamW optimizer with a maximum learning rate of 1.4351 × 10^−4^, a one-cycle learning-rate schedule, a batch size of 4,096 and 10 epochs. Hyperparameters were selected using Optuna^64^ to maximize the macro F1 score on the donor-held-out folds with raw cell-state embeddings as input. The selected settings were then held fixed across all input representations and embedding models. Using these settings, a final classifier was trained on cytokine-treated cells from all donors in Parse-PBMC, with PBS cells used to construct the control references. This final training used no held-out split, and the resulting classifier was used for downstream inference on external query datasets.

For inference, the input representation was constructed according to the availability of donor-matched controls in the query dataset.

#### Paired datasets

When donor-matched controls were available, inputs were constructed using the same formulation as in training. Control centroids were computed separately for each donor and annotated cell type, and the pooled control centroid for each cell type was calculated from the control cells of all donors in the query dataset. These centroids were used to construct the donor-adjusted receiver state and matched deviation, which were concatenated. For the macrophage dataset, which comprised a single cell type, matched control centroids were computed per donor. Cells lacking a cell-type annotation and donor-cell-type groups without control cells were excluded from paired reference.

#### Unpaired datasets

When disease and control samples originated from different donors, observed donor-matched control centroids were unavailable for the disease samples. We therefore used one of two alternative control references.

For the pooled control reference, the centroid 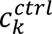 was computed from all control cells of annotated cell type *k* in the query dataset. Because this reference does not provide an estimate of the donor-specific baseline offset, the input concatenated the raw cell-state embedding with its deviation from the pooled control centroid:

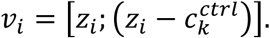

For the counterfactual control reference, CellOT^6^ was trained onto the control cells of the query dataset and used to generate model-imputed control states for non-control cells. These states were averaged within each donor and annotated cell type to obtain the counterfactual control centroid 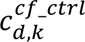 The pooled control centroid 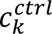 was calculated from the observed control cells as described above. Substituting the counterfactual centroid for the observed donor-matched control centroid preserved the paired input formulation:

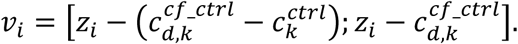

Cells in the control group were not transported and were represented using their own observed donor- and cell-type-matched control centroids. Details of counterfactual generation and the CellOT settings are provided in Supplementary Note 2.

### Statistical analysis and effect size estimation

Predicted cytokine probabilities for each cell *i* and cytokine *c* were clipped to the interval [10^−6^, 1 − 10^−6^] to avoid infinite values during logit transformation. The per-cell cytokine response score was then defined as

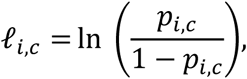

where *p_i_*_,*c*_ denotes the clipped probability. To avoid treating individual cells as independent biological replicates, statistical inference was performed using donor-level summary scores. For each donor *d*, annotated cell type *k* and condition *cond* (control or stimulated/disease), the median response score was calculated as

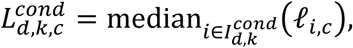

where 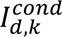 denotes the set of cells from donor *d*, cell type *k* and condition *cond*. For analyses pooling all cell types, the median was calculated across all cells within each donor-condition group, without stratification by cell type.

Effect size (Δ logit) was used as the primary measure of condition-associated cytokine response and was defined as the difference between the stimulated (or disease) and control groups in the mean of donor-level median scores. For paired control-stimulated comparisons, increases in response scores were tested using one-sided paired Student’s t-tests on the donor-level summaries, including only donors represented in both conditions. For each analyzed cell population and condition comparison, *P* values were adjusted across the 90 cytokines using the Benjamini– Hochberg false discovery rate (FDR) procedure.

### Correlation analysis with disease severity

Clinical severity categories were encoded as ordinal scores (0, healthy; 1, mild; 2, severe; 3, critical COVID-19). Analyses were performed at the donor level using median cytokine response scores calculated across all cells of each donor for pooled analyses or within each donor–cell-type group for the cell-type-resolved analyses. Spearman’s rank correlation was calculated between ordinal severity and donor-level median response scores for each cytokine in pooled analyses and for each cell-type– cytokine pair in cell-type-resolved analyses. Simple linear regression of the donor-level median response scores on ordinal severity was additionally used to estimate the direction and magnitude of score changes on the assigned severity scale. For each analyzed cell population, *P* values from the Spearman correlations were adjusted across the 90 cytokines using the Benjamini–Hochberg FDR procedure. Cytokine responses were considered to increase significantly with severity when the FDR was below 0.05 and the regression slope was positive, and were ranked by Spearman’s correlation coefficient.

Plasma protein concentrations were also assessed for association with ordinal severity using Spearman’s rank correlation. Among donors with both scRNA-seq and plasma measurements, we further calculated Spearman correlations between plasma cytokine concentrations and the corresponding CYTAP response scores. These scores were inferred using counterfactual control references and summarized as medians across all cells of each donor. Donors with plasma concentrations below the assay’s lower limit of detection were excluded from these plasma-based correlation analyses. Receptor gene expression was summarized as the mean log-normalized expression within the same donor–cell-type groups used for the cell-type-resolved response analyses.

### Cytokine response signature scores

Cytokine response signature scores were calculated from log-normalized expression using the score_genes function in Scanpy (v1.11.5). Gene sets were retrieved using GSEApy^65^ and restricted to genes detected in each dataset. The IFN-β and IL-4 signature scores were calculated using the Gene Ontology gene sets “response to interferon-β” (GO:0035456) and “response to interleukin-4” (GO:0070670), respectively. The IFN-γ, IL-6 and TNF-α signature scores were calculated using the MSigDB Hallmark gene sets^66^ “Interferon gamma response”, “IL-6/JAK/STAT3 signaling” and “TNF-α signaling via NF-kB”, respectively.

### Variance partitioning of input representations

To characterize the source of variation retained by each input representation, we computed centroids of cytokine-treated cells for each donor–cell-type–treatment combination and performed variance partitioning in the full centroid feature space, without principal component truncation. A fixed-effects model included main effects for cell type, donor and treatment, together with donor × cell type and treatment × cell type interactions. The contribution of each model term was quantified by hierarchical partitioning^67,68^, and expressed as a percentage of the total sum of squares for the corresponding representation. Variation not explained by the fitted model was assigned to the residual component. Details of the design matrix, partitioning procedure and implementation are provided in Supplementary Note 3.

### Ablation of input representations

To assess the predictive contribution of each component of the composite representation and compare embedding-based inputs with a gene-expression baseline, we performed four-fold donor-held-out cross-validation on the Parse-PBMC dataset. Separate classifiers were trained on five input representations: log-normalized expression of 2,058 HVGs (matching the dimensionality of the raw embedding), raw embeddings *z_i_*, donor-adjusted receiver states *~z_i_*, matched deviations *δ_i_*, and composite representations *v_i_*. The MLP hidden-layer architecture and training hyperparameters were held constant across representations, with the input-layer dimensionality adjusted accordingly (see Model training and inference).

HVGs were selected once from the full Parse-PBMC dataset before cross-validation using the highly_variable_genes function in Scanpy with batch_key set to donor. The same gene set was used in all folds. Thus, held-out donors contributed to HVG selection but not to classifier fitting. In each fold, three of the 12 donors were reserved for validation, and the remaining nine were used for training. All cells from the held-out donors, including their PBS controls, were excluded from classifier training, and pooled control centroids were computed using PBS control cells from the training donors only. For paired validation, donor- and cell-type-matched control centroids were calculated from the held-out donors’ own PBS control cells and used to construct their input representations. Prediction performance was evaluated overall and within each cell type using macro F1 score and macro-averaged one-vs-rest AUROC.

To assess transfer to an independent dataset, classifiers trained on the full Parse-PBMC dataset with each of the five input representations were applied to the macrophage dataset using the corresponding input construction procedures. For the HVG input, expression values for HVGs absent from the query dataset were set to zero.

### Comparison of pretrained cell embeddings

To assess the effect of pretrained embedding choice on cytokine-response prediction, we repeated the donor-held-out evaluation described above using raw cell-state embeddings from five single-cell foundation models: State SE, scFoundation, Geneformer v2, scGPT and UCE^1–4,36^. Embeddings were extracted using the default settings of each model. For scFoundation, 3,072-dimensional embeddings were generated using the official get_embedding.py script in single-cell cell-embedding mode. For Geneformer v2 104M, 768-dimensional embeddings were extracted from the CLS token representation in the penultimate layer. For UCE, 1,280-dimensional embeddings were generated using the four-layer model and eval_single_anndata.py with the species set to human. For scGPT, 512-dimensional embeddings were generated using the pretrained human model. Raw counts were supplied to scFoundation, Geneformer v2 and UCE, whereas log-normalized counts were supplied to scGPT. Gene identifiers were mapped to each model’s vocabulary before embedding extraction. The classifier input-layer dimensionality was adjusted to match each embedding, while the remaining layers, loss function, optimization settings and donor folds were held constant across models.

To examine whether the input transformation strategy extended to another embedding model, we additionally evaluated scFoundation using four input representations. Control centroids were computed in the scFoundation embedding space, and the representations were constructed and evaluated using the same procedures described above.

### Evaluation of control references for unpaired inference

To evaluate alternative control references for inference without donor-matched controls, we used the Parse-PBMC dataset, in which observed donor- and cell-type-matched PBS control centroids provided a benchmark reference. Classifiers trained in the four donor-held-out folds were applied without retraining to cytokine-treated cells from the corresponding held-out donors (see Ablation of input representations). Three reference types were compared: the observed donor- and cell-type-matched PBS control centroid 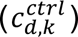, a cell-type-matched pooled control centroid 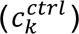, and a counterfactual control centroid obtained by averaging CellOT-generated control states within each held-out donor and cell type 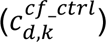.

To assess whether the relative performance of pooled and counterfactual references depended on the control population used, we evaluated each alternative using PBS-treated cells from either the held-out donors or the training donors as the control population. Pooled centroids were calculated separately for each annotated cell type within the designated control population. For CellOT, two maps were fitted in each fold using cytokine-treated cells from the held-out donors as the source population, with cells from all cytokine treatments pooled into a single group and donor labels omitted from transport fitting. One map used PBS-treated cells from the held-out donors as the target population, whereas the other used PBS-treated cells from the training donors. Thus, two CellOT settings differed in the donor origin of the target control population, not the source population. Details of transport fitting and counterfactual generation are provided in Supplementary Note 2.

Matched and counterfactual references were used to construct both donor-adjusted receiver state and matched deviations. For the pooled-reference, the composite input concatenated the raw embedding with its deviation from the pooled control centroid, as described for unpaired query inference (see Model training and inference). Each reference-substituted input was evaluated using the classifier trained on the corresponding input representation, with classifier parameters held fixed. AUROC and macro F1 score were computed over all held-out cells and separately within each cell type, and averaged across the four folds.

In a separate analysis, we performed variance partitioning on nine representations as described above: the raw embedding; the donor-adjusted state, deviation and composite representations constructed using either matched or counterfactual references; and the deviation and composite representations constructed using the pooled reference. For this analysis, pooled control centroids were computed from PBS-treated cells of all 12 donors. Counterfactual centroids were obtained using a CellOT map fitted from the cytokine-treated cells to the PBS-treated cells of all 12 donors.

### Data collection and pre-processing

The Parse-PBMC dataset profiles approximately 9.7 million peripheral blood mononuclear cells (PBMCs) from 12 donors after 24 h of in vitro stimulation with each of 90 cytokines or PBS. The quality-controlled data were downloaded and analyzed with the provided cell type annotations.

Count matrices of IFN-β-treated PBMC data^37^ were downloaded from the Gene Expression Omnibus (GEO; GSE96583). This dataset profiles PBMCs from eight patients with systemic lupus erythematosus (SLE) after 6 h of in vitro stimulation with IFN-β, along with an untreated control from each patient. Only cells annotated as singlets with valid cell type annotations (excluding ‘NA’) were retained for analysis.

Single-cell RNA sequencing (scRNA-seq) data of cytokine-treated blood-derived macrophages^39^ were downloaded from GEO (GSE168710). The count matrix was log-normalized for signature scoring and for the transcriptome-based embedding visualization. Fibroblast co-culture conditions were excluded because macrophages under those conditions are exposed to fibroblast-derived signals in addition to the applied cytokine.

The COMBAT dataset^45^ was obtained from the Chan Zuckerberg CELLxGENE data portal^69^. Red blood cells and annotated doublets were removed. Cell types comprising fewer than 500 cells (iNKT, RET, and Mast cells) as well as three samples with 500 or fewer cells were also excluded. For donors with multiple or longitudinal samples, a single representative sample per donor was retained, prioritizing the highest disease severity followed by the longest duration since symptom onset. Analysis was restricted to healthy volunteers (HV) and to COVID-19 patients with mild (COVID_MILD), severe (COVID_SEV) or critical (COVID_CRIT) disease. The correlation with severity was computed on all retained cells, whereas the cell-type-resolved comparison additionally excluded donor-by-cell-type groups containing 100 or fewer cells.

Plasma protein concentrations for the same cohort were obtained from the Luminex immunoassay data deposited with the COMBAT study^45^, which profiles 51 proteins across plasma and serum samples. Only plasma samples from healthy volunteers and from COVID-19 patients (mild, severe or critical) were analyzed. Values recorded as zero, representing concentrations below the lower limit of detection (LOD) that could not be quantified, were treated as missing values. A single sample per donor was selected using the same criteria applied to the single-cell data, and Luminex plasma samples were matched to scRNA-seq samples by donor and time point.

scRNA-seq data of bone marrow hematopoietic cells from healthy donors sampled before and after G-CSF treatment^41^ were downloaded from GEO (GSE193138), comprising four samples from two donors. Cells were filtered using the thresholds reported in the original study (at least 1,000 UMI counts, 300 detected genes and <10% mitochondrial reads). Log-normalized counts in Seurat (v4.0.3) ^70^ were used for principal component analysis on 2,000 highly variable genes, and donor effects were corrected using Harmony (v1.2.0) ^71^. Clusters were annotated as hematopoietic stem and progenitor cell (HSPC), myeloid, erythroid–megakaryocyte, T/NK, or B lineage cells based on the expression of CD34, CD33, GATA1, CD7, CD19 and *CD79A*, following the original study. The myeloid lineage population was used for comparing cytokine predictions before and after treatment.

scRNA-seq data of peripheral T cells from patients with glioblastoma or colorectal cancer sampled before and after treatment with rhIL-7-hyFc^42^ were obtained from Zenodo. Log-normalized counts were used for principal component analysis on 2,000 highly variable genes, and patient effects were corrected using Harmony. Clusters lacking T cell marker (*CD3D*) expression were removed, and the remaining cells were re-clustered and annotated on the basis of CD4, CD8A and CD8B expression. CD4 and CD8 T cells were analyzed separately.

### Dimensionality reduction and visualization

UMAP embeddings shown for the query datasets were computed from the State cell-state embeddings with the neighbors and umap functions in Scanpy at default settings. For the macrophage dataset, an additional UMAP was computed from log-normalized expression using 2,000 HVGs and the first 15 principal components, to show the position of TNF-α-treated cells relative to the untreated control. Principal component plots of the donor–cell type–treatment centroids used for variance partitioning were computed separately for each input representation.

### Benchmarking against existing methods

CYTAP was compared with NicheNet, huCIRA, SecAct, CytoSig, IREA and ContactTracing on the Parse-PBMC dataset. Responsive conditions were defined as cell-type–cytokine combinations with at least 10 differentially expressed genes (|log2FC| ≥ 0.25 and adjusted *P* < 0.05) relative to PBS-treated cells of the same cell type, based on the differential expression results provided with the dataset. This yielded 1,326 conditions spanning 87 of the 90 cytokines; C3a, IL-18 and TGF-β1 did not meet this criterion in any cell type.

For each condition, candidate signals were ranked by comparing cytokine-treated and PBS-treated cells of the same cell type, and the rank assigned to the applied cytokine was recorded. CYTAP ranked all 90 cytokines by effect size (Δ logit), with each donor’s cells scored using the classifier from the cross-validation fold in which that donor was held out (see Ablation of input representations). The other methods were run using their released models and reference resources; the huCIRA dictionary was derived from the same Parse-PBMC dataset. Cytokines were mapped to each method’s ligands, receptors or signatures using the cytokine–ligand and cytokine–receptor mapping table (see Data availability). When a cytokine mapped to multiple candidates, its best rank among those candidates was used.

For NicheNet, SecAct and CytoSig, candidate pools comprised ligands or signatures mapped to the 90 CYTAP cytokines. huCIRA and IREA ranked the cytokines represented in their dictionaries for the corresponding cell type. ContactTracing ranked the 87 benchmark cytokines for which at least one receptor was tested in the relevant cell type. A condition was considered evaluable for a method only if the applied cytokine was represented in its candidate pool and the method returned a ranking. Conditions that did not meet these requirements were excluded from that method’s evaluation rather than counted as misses.

Performance was summarized as top-k recall, defined as the fraction of evaluable conditions in which the applied cytokine ranked within the top k candidates. For a condition with n candidates, the chance expectation was calculated as *min*(*k*, *n*)/*n* and then averaged over the evaluable conditions of each method. Top-5 recall above chance was calculated as (*recall*_5_ − *random*_5_)/(1 − *random*_5_), where *recall*_5_ is the observed top-5 recall and *random*_5_ is the mean chance expectation. This measure is 0 at the chance expectation and 1 for perfect recovery. Method-specific settings are provided in Supplementary Note 4.

For in vivo validation, the same methods were applied to the G-CSF-treated bone marrow and rhIL-7-hyFc-treated peripheral T cell datasets. Post-treatment cells were compared with pre-treatment cells in the myeloid lineage for G-CSF and separately in CD4+ and CD8+ T cells for rhIL-7-hyFc. Methods were compared by the rank assigned to the administered cytokine, applying each method’s default significance filter where applicable. NicheNet provided ligand rankings without significance filtering. The granulocyte reference for huCIRA and monocyte reference for IREA were used for the myeloid population. To assess the effect of candidate-pool composition, rankings were additionally evaluated after restricting candidates to ligands or receptors corresponding to the 90 CYTAP cytokines; these restricted-pool results are denoted MAP. In both in vivo datasets, the NicheNet background comprised all genes detected in the receiver population, without the expression-percentage cutoff used in the Parse-PBMC benchmark. CYTAP predictions were filtered for significance in the IL-7 dataset. Given the limited number of donors in the G-CSF dataset (n = 2), CYTAP cytokines were ranked by effect size across all 90 candidates without significance filtering.

Administered cytokines that received no rank were assigned to one of three categories: not called, when absent from the method’s significant calls, including cases in which SecAct inferred a negative activity change; not in vocabulary, when absent from the method’s reference, as for IL-7 in CytoSig; or test not possible, when fewer than 10 genes overlapped the reference gene sets, as for huCIRA in the G-CSF dataset. Methods were also characterized by whether they provide single-cell resolution, operate without a preselected receiver cell type and rank candidates without filtering on ligand or receptor expression (Fig. 5f). NicheNet was classified as partially meeting the last criterion because its expression cutoff is applied by default but can be omitted. Scupa was evaluated separately because it reports predefined polarization states rather than individual cytokine activities (Supplementary Note 4).

## Statistics and reproducibility

No statistical method was used to predetermine sample size. All analyses used previously published datasets, with the cell and sample exclusions described above. Randomization and blinding were not applicable because no new experiments were performed. Donor-held-out cross-validation folds were fixed in advance and shared across input representations and embedding models. Predictions were evaluated across the four folds and in five independent datasets. Statistical tests and the unit of analysis are given in the figure legends and in Statistical analysis and effect size estimation.

## Data availability

The Parse-PBMC dataset is available from [https://www.parsebiosciences.com/datasets/10-million-human-pbmcs-in-a-single-experiment/]. The IFN-β-treated PBMC and cytokine-treated macrophage datasets are available from the Gene Expression Omnibus under accession numbers GSE96583 and GSE168710, respectively. The COMBAT dataset is available from the CZ CELLxGENE Discover portal [https://cellxgene.cziscience.com/collections/8f126edf-5405-4731-8374-b5ce11f53e82]. The G-CSF- and IL-7-treated datasets used for in vivo validation are available under accession number GSE193138 and at Zenodo [https://zenodo.org/records/17958065] respectively. The COMBAT Luminex plasma measurements are available from the Zenodo record of the COMBAT study [https://zenodo.org/records/6120249]. State embeddings and cell metadata of the query datasets, CYTAP predictions, the trained models, the result tables underlying the figures and the cytokine–ligand and cytokine–receptor tables used for benchmarking are available at Zenodo [https://doi.org/10.5281/zenodo.24012391].

## Code availability

The CYTAP Python package, including a tutorial and the pretrained model, is available at GitHub [https://github.com/CB-postech/CYTAP]. Code to reproduce the analyses in this study is available at GitHub [https://github.com/CB-postech/CYTAP-paper].

## Acknowledgements

This work was supported by grants from the National Research Foundation of Korea (Grant No.: RS-2023-NR077247, RS-2023-00221112, RS-2024-00426031, RS-2025-02216523, RS-2026-25511199, and RS-2026-25616628 to J.K.K., RS-2025-25431391 to Y.L.J.). Claude Opus 5 and Opus 5.5 were used to improve and organize the analysis code and to improve the clarity of the manuscript. All code and text were reviewed and verified by the authors.

## Author contributions

J.K.K conceived and supervised the study. Y.L.J performed the bioinformatic and data analyses. Y.L.J and J.K.K. wrote the manuscript.

## Extended Data Figures and legends

**Extended Data Figure 1.**
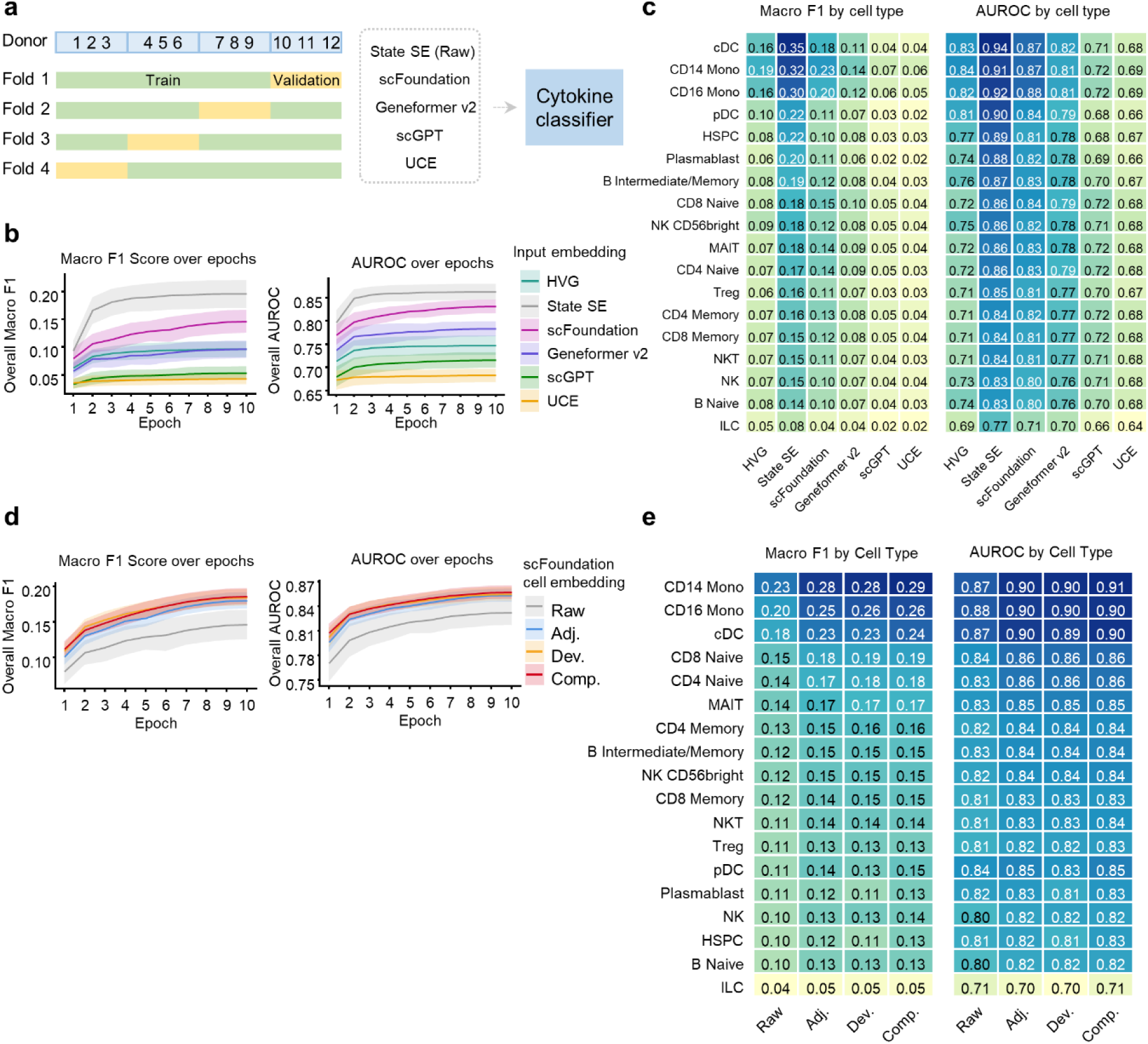
Choice of cell-state embedding and generalization of the input representation. a, Schematic of the experiment. The donor-level four-fold split and classifier of Fig. 2a were used, with cell embeddings from five single-cell foundation models as input: State SE, scFoundation, Geneformer v2, scGPT and UCE. The input layer was sized to each model’s embedding dimension, while the remaining layers were identical across models. The HVG input is included as a baseline in b and c. **b,** Macro F1 score and AUROC on the held-out validation donors over 10 training epochs. Lines, mean across the four folds. Shading, s.d. across the four folds. **c,** Heatmap of per-cell-type macro F1 and AUROC at epoch 10, averaged across the four folds. Cell types are ordered by their score under State SE. **d,** Macro F1 score and AUROC on the held-out validation donors over 10 training epochs for classifiers trained on the scFoundation embedding, used directly as the raw cell state (Raw) or transformed into the Adj., Dev. and Comp. representations defined in Fig. 2a. Lines, mean across the four folds. Shading, s.d. across the four folds. **e,** Heatmaps of per-cell-type macro F1 and AUROC at epoch 10 for the same four inputs, averaged across the four folds.

**Extended Data Figure 2.**
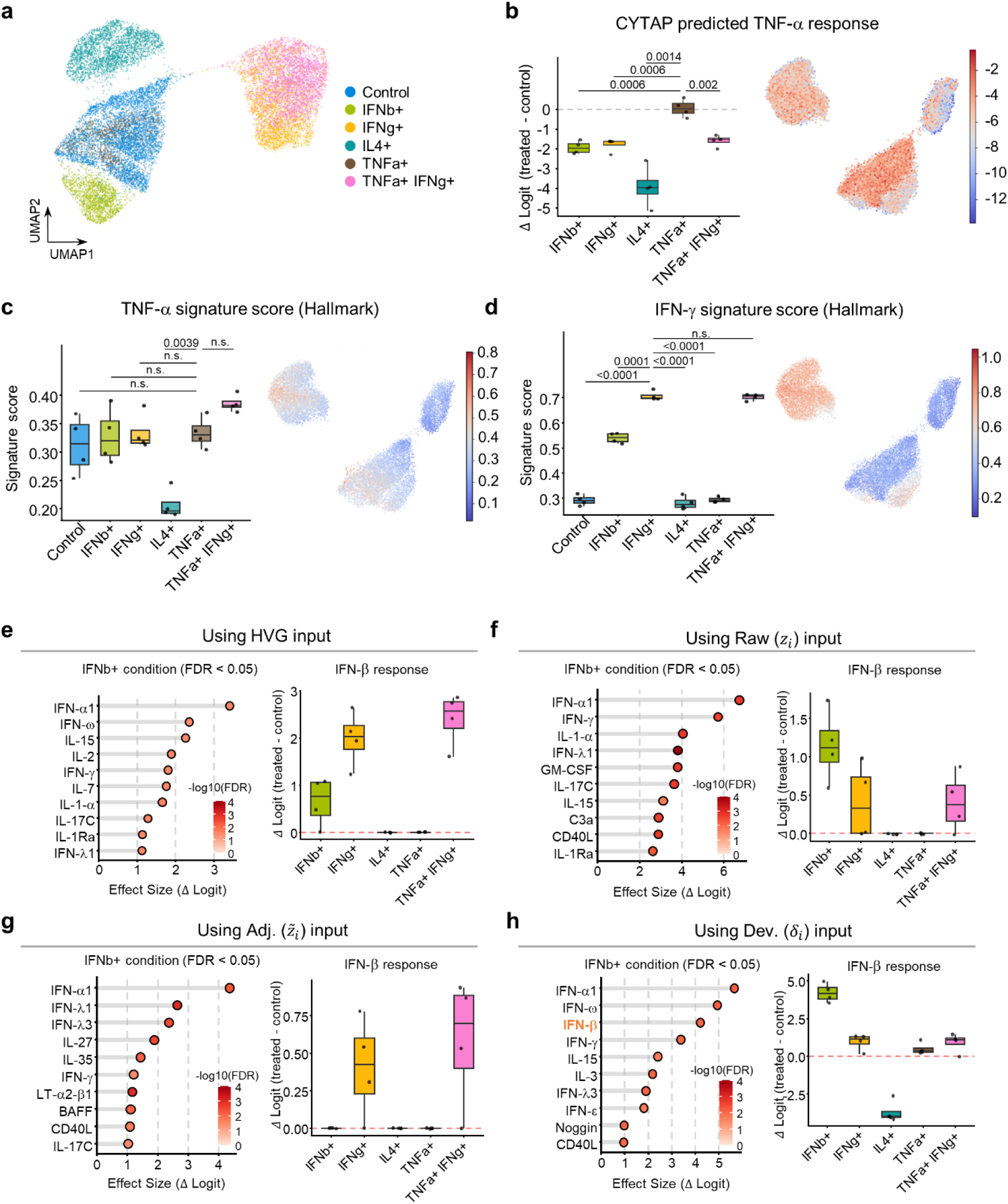
Additional conditions and input representations for the cytokine-treated macrophage dataset. a, UMAP embedding computed from the 2,000 most highly variable genes, colored by condition. **b,** CYTAP-predicted response to TNF-α. Box plots (left) show the response in each treatment condition, expressed as the change in logit relative to each donor’s control (Δ logit). Each dot is one donor. Center line, median; box, interquartile range; whiskers, 1.5× interquartile range. UMAP embeddings (right) are colored by the predicted logit score. *P* values are from one-sided paired t-tests across four donors (n.s., *P* > 0.05). **c,d,** Response signature scores by condition for TNF-α (c; Hallmark TNF-α signaling via NF-kB) and IFN-γ (d; Hallmark interferon gamma response), shown as box plots (left) and on UMAP embeddings (right). *P* values are from one-sided paired t-tests across four donors (n.s., *P* > 0.05). **e–h,** Cytokine predictions for IFN-β-treated versus control macrophages, from classifiers trained on the Parse-PBMC dataset with HVG (e), Raw (f), Adj. (g) or Dev. (h) as the input representation (Fig. 2a). Lollipop plots (left) show the top 10 cytokines with the largest increase in predicted response (Δ logit; FDR < 0.05; one-sided paired t-test with Benjamini–Hochberg correction, n = 4 donors), with IFN-β shown in bold orange. Box plots (right) show the predicted IFN-β response by condition. Each dot is one donor. The corresponding results for the composite representation are shown in Fig. 4b,e.

**Extended Data Figure 3.**
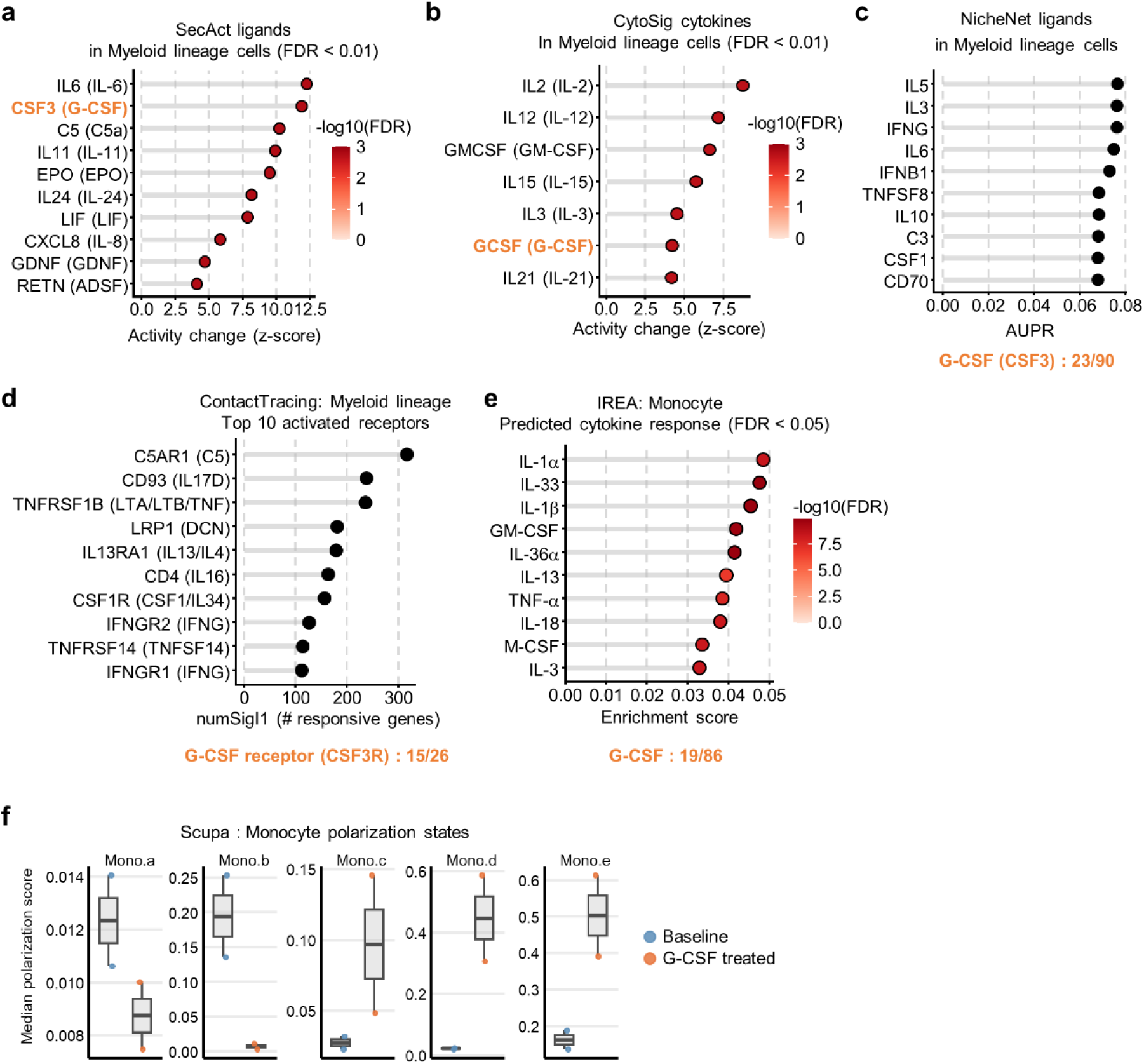
Cytokine inference by other methods in the G-CSF-treated bone marrow dataset. a–e, Lollipop plots of the candidates predicted by each method in myeloid lineage cells after versus before G-CSF treatment. IREA (e) was run with monocytes as the proxy cell type. G-CSF is shown in bold orange. Where it is not among the top 10, its rank is given below the plot. FDR thresholds for SecAct (a) and CytoSig (b) follow each method’s default. SecAct (a), NicheNet (c) and ContactTracing (d) candidates were restricted to ligands or receptors of the 90 CYTAP cytokines. **a,** Top 10 ligands predicted by SecAct (FDR < 0.01), labeled with the corresponding cytokine. **b,** All seven cytokines predicted by CytoSig (FDR < 0.01), labeled with the CytoSig name followed by the CYTAP name. **c,** Top 10 ligands predicted by NicheNet. **d,** Top 10 activated receptors predicted by ContactTracing, labeled with the corresponding ligand. **e,** Top 10 cytokines with enriched responses predicted by IREA (FDR < 0.05). Cytokine names follow IREA’s nomenclature. **f,** Box plots of the median polarization score per donor for the five monocyte polarization states predicted by Scupa (Mono.a–Mono.e), shown for the baseline and G-CSF-treated conditions. Center line, median; box, interquartile range; whiskers, 1.5× interquartile range.

**Extended Data Figure 4.**
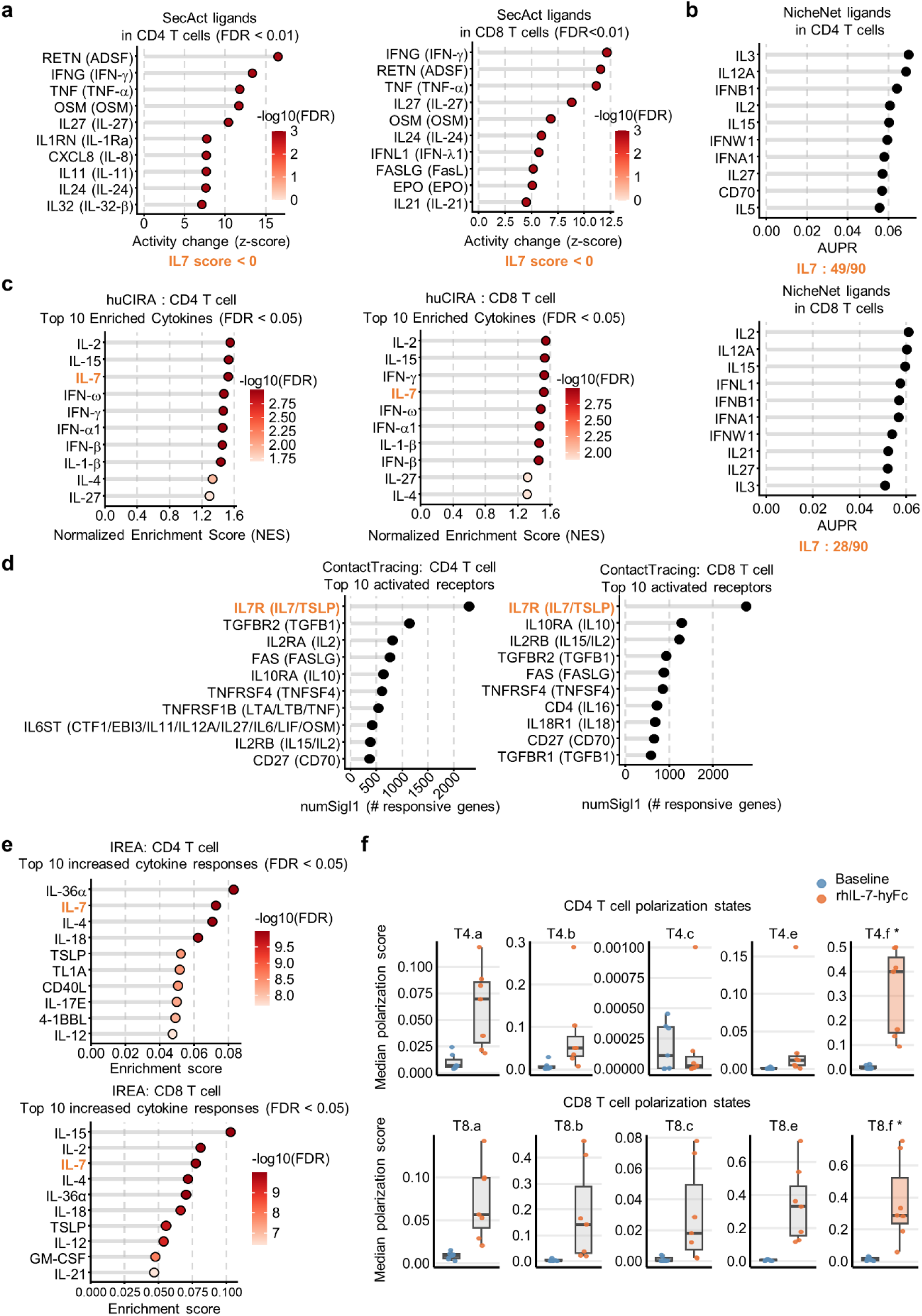
Cytokine inference by other methods in the rhIL-7-hyFc-treated peripheral T cell dataset. a–e, Lollipop plots of the candidates predicted by each method in CD4 or CD8 T cells after versus before rhIL-7-hyFc treatment. IL-7 is shown in bold orange. Where it is not among the top 10, its rank is given below the plot. The FDR threshold for SecAct (a) follows the method’s default. SecAct (a), NicheNet (b) and ContactTracing (d) candidates were restricted to ligands or receptors of the 90 CYTAP cytokines. **a,** Top 10 ligands predicted by SecAct (FDR < 0.01), labeled with the corresponding cytokine. IL-7 showed a negative activity change and was therefore not called, as noted below the plot in place of its rank. **b,** Top 10 ligands predicted by NicheNet. **c,** Top 10 cytokines predicted by huCIRA (FDR < 0.05). **d,** Top 10 activated receptors predicted by ContactTracing, labeled with the corresponding ligand. **e,** Top 10 cytokines with enriched responses predicted by IREA (FDR < 0.05). Cytokine names follow IREA’s nomenclature. **f,** Box plots of the median polarization score per patient for the five CD4 and five CD8 T cell polarization states predicted by Scupa (T4 and T8 states a, b, c, e and f), shown for the baseline and rhIL-7-hyFc-treated conditions. IL-7 is one of the cytokines that induce the T4.f and T8.f states, which are marked with a colored box and an asterisk. Center line, median; box, interquartile range; whiskers, 1.5× interquartile range.

**Extended Data Figure 5.**
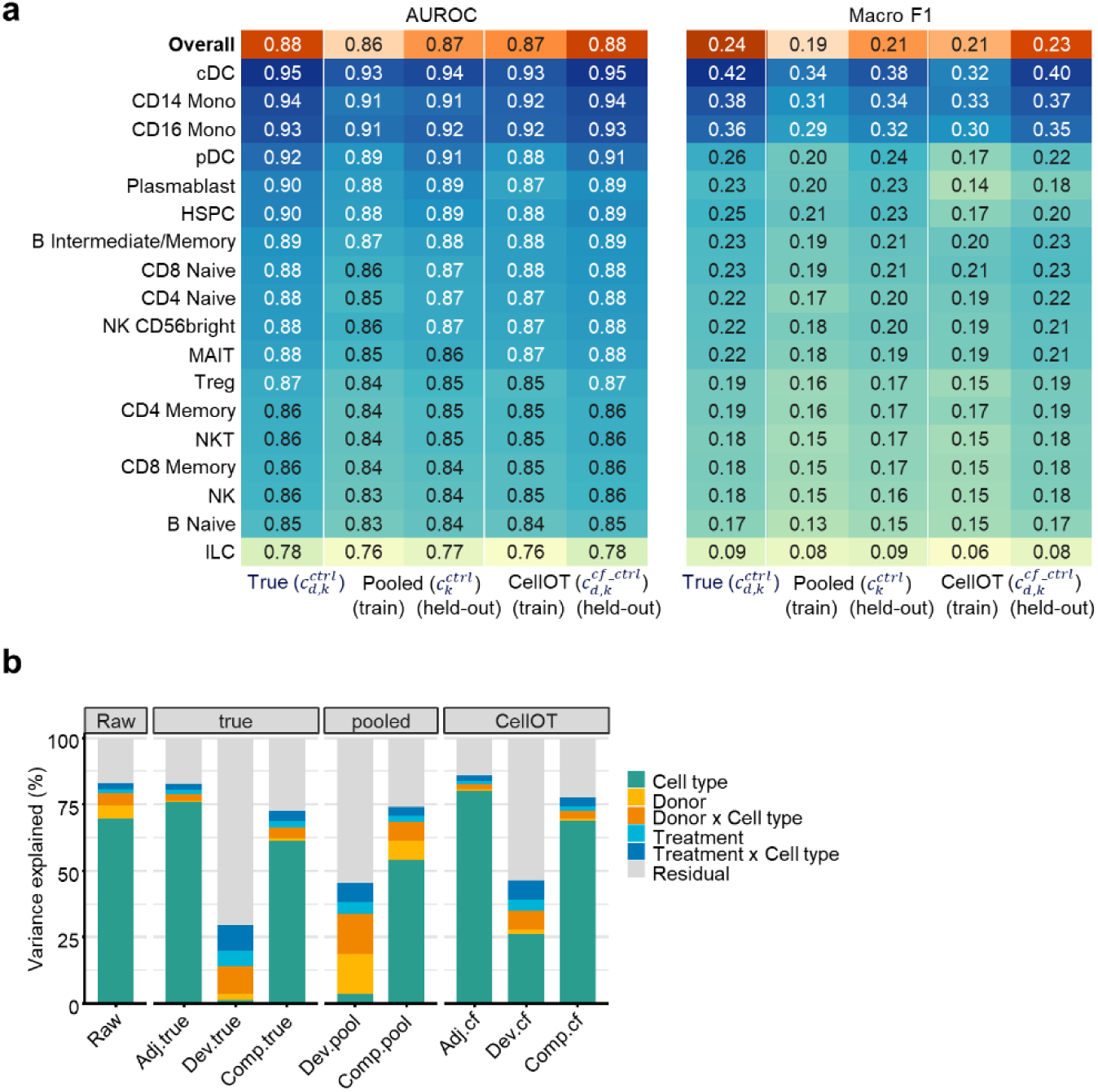
Choice of control reference for unpaired inference. a, Comparison of control references for unpaired inference. The cytokine classifier trained on the composite representation with the donor-level four-fold split of Parse-PBMC was applied to cells from the held-out donors, with the control reference set to the true donor- and cell-type-matched centroid 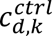, a centroid of the same cell type pooled across donors 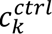, or a CellOT-generated counterfactual control centroid 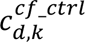. Labels in parentheses indicate whether the PBS controls used to compute the pooled centroid or define the CellOT target distribution came from the training donors (train) or the held-out donors (held-out). In both CellOT settings, the source population comprised cytokine-treated cells from the held-out donors. Heatmaps show AUROC and macro F1 computed over all held-out cells (Overall) and within each cell type, averaged across the four folds. **b,** Hierarchical partitioning of variance for centroids of each donor, cell type and cytokine, built from the nine representations defined below. The pooled reference does not define an adjusted state, so no Adj.pool representation exists, and Comp.pool combines Raw with Dev.pool. Bars show the contribution of cell type, donor, donor × cell type, treatment, treatment × cell type and residual variation, each expressed as a percentage of that representation’s own total variance. The Raw and true representations are those shown in Fig. 2e, where Adj.true, Dev.true and Comp.true appear as Adj., Dev. and Comp. For b, pooled centroids were computed from all 12 donors, and counterfactual centroids were obtained from a CellOT map fitted from their cytokine-treated cells to their PBS controls. Representations: Raw, *z*; Adj.true, *z* − 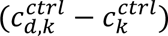; Dev.true, 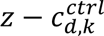 Comp.true, 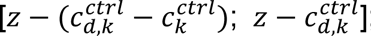; Dev.pool, 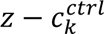; Comp.pool, 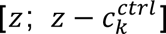; Adj.cf, 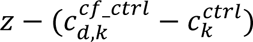; Dev.cf, 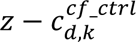; Comp.cf, 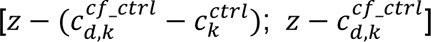.

**Extended Data Figure 6.**
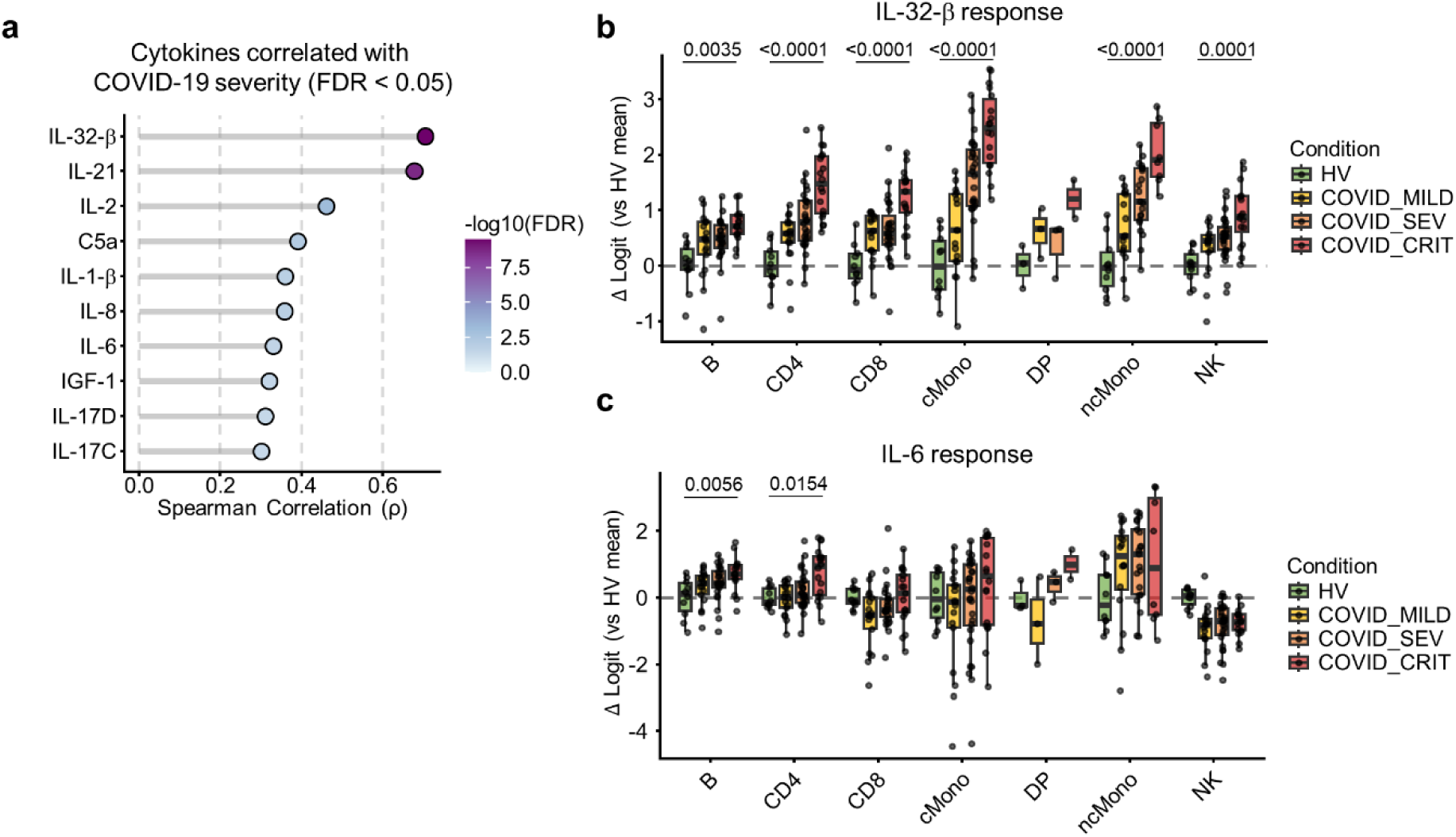
Severity-associated cytokine responses in the COMBAT COVID-19 PBMC cohort, inferred without CellOT. a–c, As in Fig. 6b–d, but with the CYTAP response inferred against a pooled control centroid instead of a CellOT-generated counterfactual control. a, Top 10 cytokines whose predicted response increases with COVID-19 severity (FDR < 0.05). b,c, Predicted response to IL-32β (b) and IL-6 (c) by cell type. FDR values are shown for cell types in which the Spearman’s correlation with severity was significant (FDR < 0.05).

**Extended Data Figure 7.**
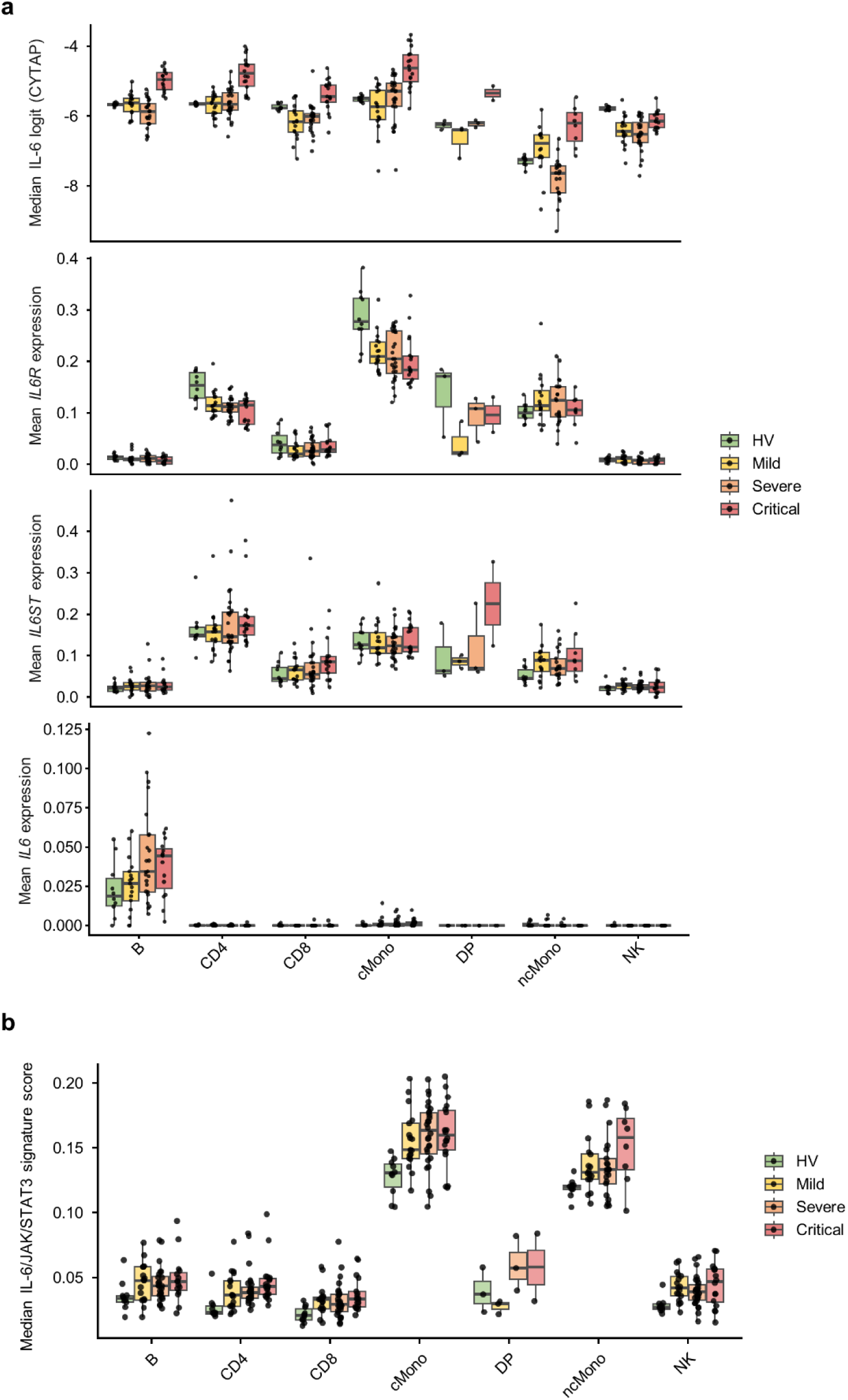
Predicted IL-6 response, expression of *IL6* and IL-6 receptor subunits, and IL-6/JAK/STAT3 signature score by cell type and severity group. a, Median CYTAP-predicted IL-6 response per donor (top), and mean log-normalized expression of *IL6R* (second row), *IL6ST* (third row) and *IL6* (bottom), for each cell type and severity group. Each dot is one donor. Cell types are restricted to those represented in all four groups. **b,** Median IL-6/JAK/STAT3 signature score per donor for the same cell types and groups. In a and b, donor–cell type combinations containing 100 or fewer cells were excluded. Box plots in a and b: center line, median; box, interquartile range; whiskers, 1.5× interquartile range.

