## Supplementary Notes for "Task-aligned foundation model representations enable single-cell cytokine response inference"

#### Supplementary Note 1. Modeling cell-state embeddings as donor-, cell-type-, and treatment-structured variation

Because cytokine responses can differ across cell types, we sought to retain receiver-cell context alongside the matched perturbation deviation. We described each cell-state embedding using an additive working model comprising a global mean, a cell-type baseline, a donor-associated baseline offset within each cell type, a cell-type-specific cytokine-response effect and a residual term. Under this model, subtracting the donor- and cell-type-matched PBS control centroid yields a matched deviation that emphasizes response-associated variation but no longer explicitly retains the cell-type control baseline. In contrast, subtracting only the estimated donor-associated baseline offset yields a donor-adjusted receiver state that retains cell-type context in a pooled cell-type reference frame. Concatenating these two vectors therefore provides the classifier with receiver-cell context alongside the deviation from the matched baseline. The assumptions and full derivation are given below.

To formalize the sources of variation represented in the latent cell-state space, we modeled each cell embedding as a combination of donor-associated baseline variation, cell-type-specific baseline state, and cytokine-induced perturbation effects. Each cell $i$ was associated with donor $d(i)$, annotated cell type $k(i)$, and treatment condition $t(i)$, where $t = 0$ denotes the PBS control condition and $t > 0$ denotes cytokine stimulation. We considered the following latent decomposition model:

$$z_{i,d(i),k(i),t(i)} = \mu + \beta_{k(i)} + \alpha_{d(i),k(i)} + \gamma_{k(i),t(i)} + \epsilon_{i,d(i),k(i),t(i)}.$$

Here, $\mu$ denotes the global mean cell state embedding, $\beta_{k(i)}$ denotes the baseline cell-type effect, $\alpha_{d(i),k(i)}$ denotes donor-associated baseline variation within cell type $k(i)$, and $\gamma_{k(i),t(i)}$ denotes the population-level cytokine response specific to cell type $k(i)$. This formulation explicitly allows cytokine responses to be cell-type specific. Thus, we did not assume a single global cytokine perturbation vector shared across all receiver cell types. Instead, the same cytokine was allowed to induce distinct response directions depending on the baseline cell state of the receiver cell type.

This model was used as a working representation model rather than as a fully identifiable generative model. Donor-specific deviations from the population-level cytokine response may exist, but such deviations cannot be reliably separated from residual variation at the donor–cell type–treatment centroid level without additional biological replication. Therefore, we did not include a separate donor-by-cell type-by-treatment response term in the primary model. Instead, donor-specific response heterogeneity, control-centroid estimation error, donor-specific substate composition, and cell-level noise were absorbed into the residual term $\epsilon_{i,d(i),k(i),t(i)}$.

For PBS control cells, we assume $\gamma_{k,0} = 0$. Therefore, the donor- and cell-type-matched PBS centroid estimates the baseline state

$$c_{d,k}^{ctrl} = \mu + \beta_k + \alpha_{d,k} + \bar{\epsilon}_{d,k}^{ctrl} \approx \mu + \beta_k + \alpha_{d,k}$$

where $\bar{\epsilon}_{d,k}^{ctrl} = \frac{1}{|C_{d,k}^{ctrl}|} \sum_{i \in C_{d,k}^{ctrl}} \epsilon_{i,d,k,0}$. The pooled PBS centroid for cell type $k$ estimates the population-level baseline

$$c_k^{ctrl} = \mu + \beta_k + \bar{\alpha}_k + \bar{\epsilon}_k^{ctrl} \approx \mu + \beta_k + \bar{\alpha}_k$$

where $\bar{\alpha}_k$ denotes the average donor-associated offset among control cells of cell type $k$ and $\bar{\epsilon}_k^{ctrl}$ denotes the corresponding average of the control residuals. Thus, the donor-specific baseline offset can be estimated as

$$\Delta_{d,k} = c_{d,k}^{ctrl} - c_k^{ctrl} = \alpha_{d,k} - \bar{\alpha}_k + (\bar{\epsilon}_{d,k}^{ctrl} - \bar{\epsilon}_k^{ctrl}) \approx \alpha_{d,k} - \bar{\alpha}_k.$$

Subtracting this offset from the original embedding gives the donor-adjusted state

$$\tilde{z}_i = z_i - \Delta_{d(i),k(i)} = \mu + \beta_{k(i)} + \bar{\alpha}_{k(i)} + \gamma_{k(i),t(i)} + \epsilon_{i,d(i),k(i),t(i)} - (\bar{\epsilon}_{d(i),k(i)}^{ctrl} - \bar{\epsilon}_{k(i)}^{ctrl}).$$

Because $\bar{\alpha}_k$ is shared by all cells of cell type $k$, it can be absorbed into $\beta_k$ without loss of generality. We therefore imposed the centering constraint $\bar{\alpha}_k = 0$, which gives

$$\tilde{z}_i = \mu + \beta_{k(i)} + \gamma_{k(i),t(i)} + \text{residual}.$$

Thus, $\tilde{z}_i$ preserves the receiver cell-type context and cytokine-induced state while reducing donor-specific baseline variation. In contrast, the matched deviation vector $\delta_i$ is approximately

$$\delta_i = \gamma_{k(i),t(i)} + \epsilon_{i,d(i),k(i),t(i)} - \bar{\epsilon}_{d(i),k(i)}^{ctrl} = \gamma_{k(i),t(i)} + \text{residual}_{i,d(i),k(i),t(i)}.$$

Thus, $\delta_i$ directly represents the cell's observed deviation from its donor- and cell-type-matched PBS baseline. The final representation $v_i$ therefore combines two complementary quantities: a donor-adjusted receiver-state context and a matched perturbation deviation. This design is important because cytokine responses are expected to be cell-type specific. The same perturbation deviation may have different biological meaning depending on the baseline receiver cell state. By including both $\tilde{z}_i$ and $\delta_i$, the model is provided with both the baseline cellular context and the direction of perturbation away from matched control. Moreover, since $\tilde{z}_i = \delta_i + c_{k(i)}^{ctrl}$, the composite representation can equivalently be viewed as providing the model with the matched perturbation residual $\delta_i$ together with the corresponding cell-type control baseline $c_{k(i)}^{ctrl}$. In this sense, $v_i = [\delta_i + c_{k(i)}^{ctrl}; \delta_i]$ implements a cell-type-conditioned cytokine response representation while reducing donor-specific baseline confounding.

The perturbation-induced deviation vector $\delta_i$ removes the donor- and cell-type-specific PBS baseline. However, because it represents only the observed deviation from the matched baseline, residual donor-structured variation in the perturbation-deviation space can become prominent. In contrast, $\tilde{z}_i = \delta_i + c_{k(i)}^{ctrl}$ expresses the same observed deviation in a pooled cell-type control reference frame. The two components are therefore expected to differ in how residual donor-structured variation is distributed, which we examined empirically using variance partitioning (Supplementary Note 3).

**Supplementary Note 2. Counterfactual control reference using CellOT**

When a query dataset contains no donor-matched control samples, a donor-matched control state can instead be generated computationally. We used CellOT, which learns a neural optimal transport map between two unpaired cell populations by parameterizing the dual potentials of the optimal transport problem with input-convex neural networks.

The cell-state embeddings were reduced to 100 principal components with a principal component analysis fitted on the cells used to train the maps, so that no information from other datasets entered the counterfactual. In the COMBAT dataset, a separate map was trained from each disease group (mild, severe and critical COVID-19) to the healthy volunteers. In the evaluation of control references, two maps were trained per fold from the cytokine-treated cells of the three held-out donors, with all cytokines pooled into one source group and without donor labels, one onto the PBS-treated cells of the held-out donors and one onto those of the nine training donors. A map trained from the cytokine-treated to the PBS-treated cells of all 12 donors was used for variance partitioning. Cells were split 8:2 into training and validation sets within each group, and each map was evaluated every 100 iterations, retaining the checkpoint with the lowest maximum mean discrepancy (MMD) between the transported and control cells of the validation sets. Each cell of a source group was transported by its map, and the result was returned to the embedding space with the inverse principal component transform to give its counterfactual control state. Cells of the control group were not transported, because these donors provide their own control cells, as in the paired setting. For each donor $d$ and cell type $k$, the counterfactual control centroid $c_{d,k}^{cf\_ctrl}$ was then defined as the mean of the counterfactual states of that donor and cell type, and used in place of the donor- and cell-type-matched PBS centroid, so that the donor-adjusted state and the matched deviation were computed exactly as in the paired formulation ($v_i = [\tilde{z}_i; \delta_i]$).

CellOT was run with the architecture and optimizer settings of its example configuration: two input-convex neural networks with four hidden layers of width 64, a convexity penalty coefficient of 1 (fnorm_penalty = 1), the Adam optimizer with a learning rate of $1 \times 10^{-4}$ and $\beta = (0.5, 0.9)$, a batch size of 256 (1,024 for Parse-PBMC) and 10 inner iterations per outer iteration. Models were trained for 10,000 iterations. The trained maps were applied using a reimplementation of the CellOT transport step that reproduces the outputs of the original implementation.

### Supplementary Note 3. Variance partitioning of the composite input representation

We compared the composite input representation $v_i = [\tilde{z}_i; \delta_i]$ with its two components and the raw embedding by variance partitioning. For each representation $r \in \{z_i, \tilde{z}_i, \delta_i, v_i\}$, we computed donor–cell type–treatment centroids of the cytokine-treated cells and performed the decomposition in the full centroid feature space, without principal component truncation, so that every representation was compared on its total variance rather than on a representation-dependent subset of it. Let $Y^{(r)}$ denote the centroid matrix for representation $r$, $\bar{Y}^{(r)}$ its grand mean matrix, and $\hat{Y}_M^{(r)}$ the matrix of fitted values under a fixed-effect model $M$. Explained variance was quantified as a sum of squares,

$$SS_M(r) = \left\|Y^{(r)} - \bar{Y}^{(r)}\right\|_F^2 - \left\|Y^{(r)} - \hat{Y}_M^{(r)}\right\|_F^2,$$

where the first term is the total sum of squares, denoted $TSS(r)$.

Variance components were defined as term blocks of a single fixed-effect design, fitted as

$$M_{full} : Y^{(r)} \sim \text{cell type} + \text{donor} + \text{donor: cell type} + \text{treatment} \\ + \text{treatment: cell type}$$

where a colon denotes the interaction contrast, so that each of the five terms corresponds to one variance component: cell type (receiver identity), donor, donor:cell type, treatment (shared cytokine response) and treatment:cell type (cell-type-specific cytokine response). All factors were encoded with sum-to-zero (deviation) contrasts, and each component corresponded to a block of columns of this single design matrix. Donor-associated baseline variation was represented by a donor main effect together with a donor×cell type term, consistent with the use of donor- and cell-type-matched PBS centroids. Together, these two terms correspond to $\alpha_{d,k}$ in the latent decomposition model above, and the treatment and treatment×cell type terms together correspond to $\gamma_{k,t}$. The donor:treatment and three-way donor:cell type:treatment terms were not included, and the residual component was defined as $TSS(r) - SS_{full}(r)$.

The independent contribution of each component was obtained by hierarchical partitioning, computed as its average marginal increase in explained sum of squares over all $2^5 = 32$ component subsets, weighted so that the result is the Shapley value of that component. The decomposition was reimplemented in Python 3.11.13 with NumPy (v2.3.5), SciPy (v1.16.3) and patsy (v1.0.2) so that it could be computed in the full centroid space. The five contributions sum to $SS_{full}(r)$, and adding the residual recovers $TSS(r)$. Contributions are reported both as absolute sums of squares in embedding units and as a percentage of $TSS(r)$. Because $v_i$ concatenates two feature blocks and multivariate least squares under a shared design matrix decomposes column-wise, both the total and the component sums of squares are additive across blocks, $SS_c(v) = SS_c(\tilde{z}) + SS_c(\delta)$. Comparison across representations was therefore based on percentage contributions rather than on absolute sums of squares.

The composite representation was considered valid if the donor and donor×cell type contributions decreased relative to the raw embedding $z_i$, the treatment and treatment×cell type contributions remained detectable, and the cell type contribution, which carries receiver cell-state context, was retained. The last criterion was included because subtracting the matched control centroid removes the explicit cell-type control baseline, which is retained in the donor-adjusted receiver state.

### Supplementary Note 4. Settings used for the benchmarked cytokine-inference methods

NicheNet was run using the human ligand–target prior (v2). For each responsive condition, background genes were defined as genes expressed in at least 5% of cells of the corresponding receiver cell type, and the gene set of interest comprised the differentially expressed genes of that condition ($|\log2FC| \geq 0.25$ and adjusted $P < 0.05$) that were also background genes, both restricted to genes present in the ligand–target prior. Ligand activities were computed with predict_ligand_activities, and candidate ligands were ranked by the corrected area under the precision–recall curve (AUPR). Conditions with fewer than 10 genes in the gene set were not scored and were therefore not evaluable.

For huCIRA, enrichment of the Human Cytokine Dictionary gene sets of the matching cell type was tested for each condition with run_one_enrichment_test on log-normalized expression, contrasting cytokine-treated with PBS-treated cells, with the default settings. Cytokines were ranked by normalized enrichment score, and a condition was evaluable only when at least 10 genes were shared with the dictionary.

For SecAct, pseudo-bulk log fold changes were computed for each cell type from raw counts (log2-transformed counts per million, treated minus control) and supplied to SecAct.activity.inference with is.differential = TRUE.

CytoSig activities were computed from the same pseudo-bulk log fold-change matrix using SecAct.activity.inference with sigMatrix = “CytoSig”, lambda = 10,000 and is.group.sig = FALSE. CytoSig signature labels were mapped to cytokines using a curated dictionary in which many-to-one family mappings, such as the type I interferons, were aggregated, yielding 33 signatures that covered 36 of the 87 responsive cytokines.

For the Parse-PBMC benchmark, the gene-set mode of IREA was reproduced offline: the upregulated differentially expressed genes of each condition were tested against the human-gene versions of the IREA cytokine signatures of the matching IREA cell type by hypergeometric enrichment, and cell types without an IREA counterpart were not evaluable. For the in vivo datasets, the IREA web tool was used in Gene Matrix mode.

ContactTracing was run without its ligand-expression filter, and each cytokine was scored by the activity of its cognate receptors (the number of response genes with an interaction FDR $< 0.05$), so that cytokines could be ranked irrespective of ligand expression. Receptors expressed in at least 2% of the cells of each cell type were tested, with the highly variable genes of that cell type and the receptors as response genes. Up to 200 cells were sampled per donor, cell type and condition, a minimum of 30 cells was required, PBS-treated cells of each cell type served as controls, and the number of detected genes and donor were included as covariates.

For the in vivo datasets, treated cells were compared with baseline cells of the same population. SecAct was run with SecAct.CCC.scRNAseq and CytoSig on the pseudo-bulk log fold change of each population, the NicheNet gene set comprised the genes differentially expressed between treated and baseline cells, and ContactTracing was applied to all receptors expressed in at least 2% of the receiver cells.

Scupa reports membership in predefined polarization states rather than activity for individual cytokines and was therefore evaluated separately, on the two in vivo datasets only. UCE embeddings were computed for each dataset with eval_single_anndata.py for human, and supplied to MeasurePolar in R with embedding set to uce. The cells sampled before treatment in each population were specified as the unpolarized reference, so that polarization is measured relative to the untreated state of the same dataset. Polarization was measured against the monocyte reference type for the myeloid lineage of the G-CSF dataset, and against the CD4 and CD8 T cell reference types for the IL-7 dataset. The resulting polarization scores were summarized as the median score per donor for each state and compared between conditions.
